# Replay builds an efficient cognitive map offline to avoid computation online

**DOI:** 10.1101/2025.01.08.632067

**Authors:** Jianxin Ou, Yukun Qu, Yue Xu, Zhibing Xiao, Tim Behrens, Yunzhe Liu

## Abstract

How do humans integrate fragmented experiences into a coherent structure that supports novel inferences? Addressing this question requires tracking learning from the very first encounters. Using magnetoencephalography, we recorded human neural activity throughout the full process – from initial learning to inference. Participants first learned one-dimensional, pairwise rank relationships that collectively formed a two-dimensional (2D) conceptual map, and then inferred unobserved relationships. During rest, offline replay integrated piecemeal memories into a coherent 2D representation, predicting the emergence of a grid-cell-like code that reflected a generalizable task schema. This schema reduced the need for effortful computations during subsequent inference. During inference, two types of on-task replay emerged: a fast replay, resembling offline replay and representing the full map, and a slow replay, focused on trial-specific details. Notably, slow replay negatively correlated with both grid-like coding and inference performance. Together, these results suggest that replay builds an efficient cognitive map offline, thereby reducing reliance on deliberate computation online.

## Introduction

Our daily experiences often appear fragmented and disconnected, yet over time we integrate these pieces into structured knowledge that can be quickly generalized. For example, we might first learn isolated facts – cheetahs are fast runners, lions are predators, zebras are prey – and later understand how these animals form an interconnected ecosystem: cheetahs chase zebras, while zebras remain vigilant to evade lions. This integrated knowledge, known as a cognitive map (Tolman, 1948), enables efficient learning and flexible inference (Behrens et al., 2018). If a new predator arrives, we can quickly predict how it might disrupt the existing balance.

To build such cognitive maps, a key process is “offline replay”. During rest, the brain spontaneously reactivates past experiences in a time-compressed manner (Foster, 2017; Foster & Wilson, 2006; Lee & Wilson, 2002; Louie & Wilson, 2001; Wilson & McNaughton, 1994), accompanied by hippocampal ripple events (Buzsáki, 2015; Buzsáki & Moser, 2013). This replay has been linked to memory consolidation, experience reorganization, and even the formation of new relational structure (Barron et al., 2020; Gupta et al., 2010; Liu et al., 2019). Recent intracranial electroencephalography (iEEG) work further shows that human hippocampal ripples during rest are positively linked to later cognitive-map representations and predict inference performance (Xiao et al., 2025). However, little is known about how the brain unifies multiple pieces of knowledge – particularly those spread across more than one feature dimension – into a coherent map representation of the task space.

In two-dimensional (2D) spaces, grid cells in the entorhinal cortex (EC) encode such maps (Hafting et al., 2005; Jacobs et al., 2013). In humans, both the EC and medial prefrontal cortex (mPFC) exhibit grid-cell-like patterns that represent physical and conceptual spaces (Constantinescu et al., 2016; Doeller et al., 2010; Park et al., 2021a). This grid-like code, acting as a schema, generalizing across different environments (Hafting et al., 2005), enables inferences in new situations (Behrens et al., 2018). It is thought that having such representation can effectively bypass slower, more deliberate computations, and support “intuitive planning” (Baram et al., 2018).

Beyond rest, replay is also found during active tasks (Eichenbaum, 2017; Yu & Frank, 2015), although its precise function remains debated (Gillespie et al., 2021; Widloski & Foster, 2022). In rodents, “on-task replay” appears also in the prefrontal cortex for spatial navigation (Euston et al., 2007; Kaefer et al., 2020; Shin et al., 2019), while in humans, it has been linked to contextual understanding (Hahamy et al., 2023; Schwartenbeck et al., 2023) and prospective planning (Liu, Mattar, et al., 2021; Wimmer et al., 2023). Yet, it remains unknown how on-task replay might incorporate grid-like representations for novel inferences, especially in a 2D conceptual space.

Here, we use magnetoencephalography (MEG) to investigate how replay supports the learning of a 2D conceptual map offline during rest, and whether this formed representation diminishes the need for slower, on-task computations during inference. Participants initially learned pairwise rank relationships in each dimension (“piecemeal” learning), then inferred novel relationships they had never explicitly encountered. We hypothesize that offline replay not only consolidates memory along individual dimensions, but also actively forms an integrated 2D map representation, thus diminishing the need for on-demand, effortful computations and enhancing inference performance.

## Results

### Task hypotheses

During both learning and inference, participants engaged in “battle games” involving 25 objects, each with distinct ranks in two independent feature dimensions (attack power and defense power). Unbeknownst to participants, these 25 objects were arranged in a five-by-five conceptual map spanning the two dimensions (Fig. 1; see also Extended Data Fig. 1). Each dimension contained five objects distributed across five rank levels, forming a 2D task space.

**Fig. 1.**
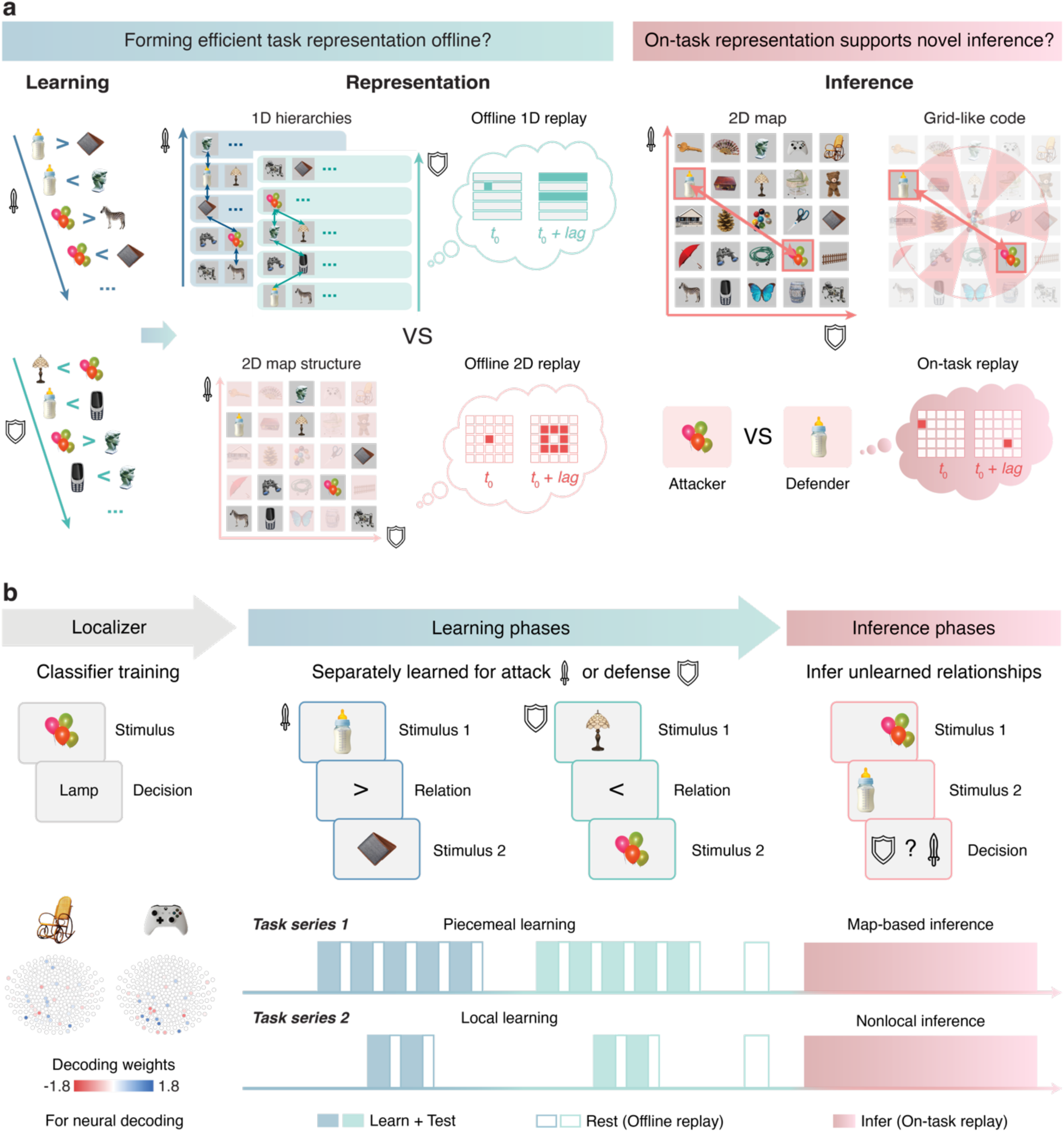
Experimental hypothesis and task design. **a**, This study investigates how offline replay during rest supports efficient learning of a 2D map representation and how that representation supports online inference of unobserved relationships. During learning, participants viewed pairwise rank relationships for 25 objects separately along each feature dimension (attack power or defense power). Based on these rank relationships, they could build either separate 1D hierarchy for each dimension, or a single, integrated 2D map, each accompanied by distinct 1D or 2D replay patterns during offline rest. During inference, we expected on-task replay of unlearned relationships alongside a grid-cell-like representation of the 2D map. **b**, In the “localizer” phase, stimulus-evoked neural activity was used to train multivariate classifiers prior to the main learning tasks, ensuring unbiased neural decoding. The experiment then comprised two learning – inference series: “piecemeal learning → map-based inference” and “local learning → nonlocal inference” (see also Extended Data Fig. 1). In both learning phases, participants observed pairwise rank relationships within a single dimension at a time, with multiple rest periods interspersed to allow spontaneous replay for offline representation learning. In both inference phases, participants inferred rank differences across dimensions, deciding whether one object’s attack power exceeded another’s defense power. We examine how the task representation is learned offline and how the formed representation interacts with on-task replay to support novel inferences.

In the learning phase, participants viewed pairwise comparisons of objects differing by one rank in a given dimension (e.g., “bottle > wallet” for attack power, Fig. 1). Because the two dimensions were introduced separately, participants could either build separate 1D hierarchy for each dimension or merge them into a unified 2D representation. We hypothesized that offline replay during rest would be crucial for forming this integrated 2D map. If replay simply consolidated single-dimensional knowledge, we would expect “1D replay” (e.g., “bottle → wallet”). Conversely, if replay supported forming an integrated task space across dimensions, we anticipated “2D replay” (e.g., object A → its neighbors on the 2D map, Fig. 1), particularly once the 2^nd^ dimension was introduced.

Following the learning phase, we examined how these representations facilitated novel inferences. During inference, participants determined which object would win based on a cued dimension (e.g., comparing *balloon*’s attack power to *bottle*’s defense power, Fig. 1a). Here, we observed a grid-cell-like “schema” representation of the 2D map and on-task replay, allowing us to investigate how offline replay, on-task replay, and grid-like coding together support inferential behavior.

### Task design and performance

Forty healthy adults participated in a full-day MEG session. The experiment consisted of a localizer phase and two main learning–inference series: “piecemeal learning → map-based inference” and “local learning → nonlocal inference.” The 1^st^ series examined how offline replay contributed to learning a 2D map representation and how the learned representation supported novel inferences, while the 2^nd^ series replicated the 1^st^ series and further tested whether the acquired representation could generalize as a schema for assimilating new objects and supporting subsequent inferences.

During the “localizer” phase, participants viewed object pictures and determined whether each picture matched a given text (Fig. 1b). Following the established pipeline (Liu et al., 2019; Liu, Mattar, et al., 2021), we used stimulus-evoked neural activity from this phase to train classifiers for unbiased replay detection.

In the 1^st^ series, during “piecemeal learning”, participants learned rank relationships between object pairs that differed by one rank. For instance, in the “attack power” dimension, they saw the “bottle” image, followed by a “>” symbol, and then the “wallet” image (Fig. 1b; Extended Data Fig. 1). The two feature dimensions (attack and defense power) were learned separately, in randomized order across participants. Each dimension’s pairs were presented twice, evenly across five blocks, ensuring that any observed performance or neural differences were not due to unequal learning experiences. After each block, participants were tested on the newly learned relationships [mean test accuracy: 0.854 ± 0.009], followed by a 1-minute rest. Completing all five blocks, they performed a pairwise test of all pairs [mean accuracy: 0.930 ± 0.012], and then took a 5-minute rest. These resting periods allowed us to detect spontaneous neural replay associated with offline representation learning.

After “piecemeal learning,” participants proceeded to the “map-based inference” phase, where they inferred unobserved relationships within each dimension or across two dimensions. Each trial presented two stimuli sequentially, followed by a decision rule signaled by “sword” (attack) and/or “shield” (defense) icons (Fig. 1b; Methods; Extended Data Fig. 1). The overall inference accuracy was 0.907 ± 0.008. We investigated whether on-task replay occurred during inferences, and how it related to behavioral performance.

In the 2^nd^ series, which mirrored the 1^st^ series, a “local learning” phase introduced four new objects into the central region of the map. Participants learned rank relationships between each new object and its neighbors in each dimension. This learning – test cycle, interspersed with rest, was identical to piecemeal learning, resulting in a memory test accuracy of 0.884 ± 0.017. Finally, in the “nonlocal inference” phase, participants inferred unobserved relationships between newly introduced objects and non-adjacent ones in the map. The procedure resembled map-based inference, except that one object was always new and the other was a non-neighbor, hence “nonlocal”. The mean inference accuracy was 0.866 ± 0.013.

### Replay detection methods

We identified spontaneous neural replay in humans using an established method, temporally delayed linear modeling (TDLM) (see Methods) (Liu, Dolan, et al., 2021; Liu et al., 2019; Liu, Mattar, et al., 2021; Nour et al., 2021). TDLM proceeds in three key steps. First, stimulus-evoked neural activity recorded during the localizer task is used to train multivariate classifiers for each object (Fig. 2a). Next, these trained classifiers are applied to the subsequent resting or on-task data, generating time series of state reactivations (Fig. 2b). Finally, a two-stage general linear model (GLM) evaluates replay transitions of interest. In the first-stage GLM, we estimate the reactivation strength for each transition pair (*β*_*transition*_) and its time-modulated component (*β*_*transition*_) at specified lags (e.g., 10, 20, 30 ms; Fig. 2c). In the second-stage GLM, we compute the replay strength (“sequenceness,” *β_replay_*) and its changes over time (“sequenceness changed over time”) by mapping these empirical transition matrices onto hypothetical replay transitions (Fig. 2d). Significance is assessed via a non-parametric permutation test, where shuffled transition matrices are entered into the second-stage GLM, and the 95^th^ percentile of the maximum permuted sequenceness distribution across time lags is used as a threshold. Any sequenceness at a specific time lag exceeding this threshold is deemed significant (Fig. 2e).

**Fig. 2.**
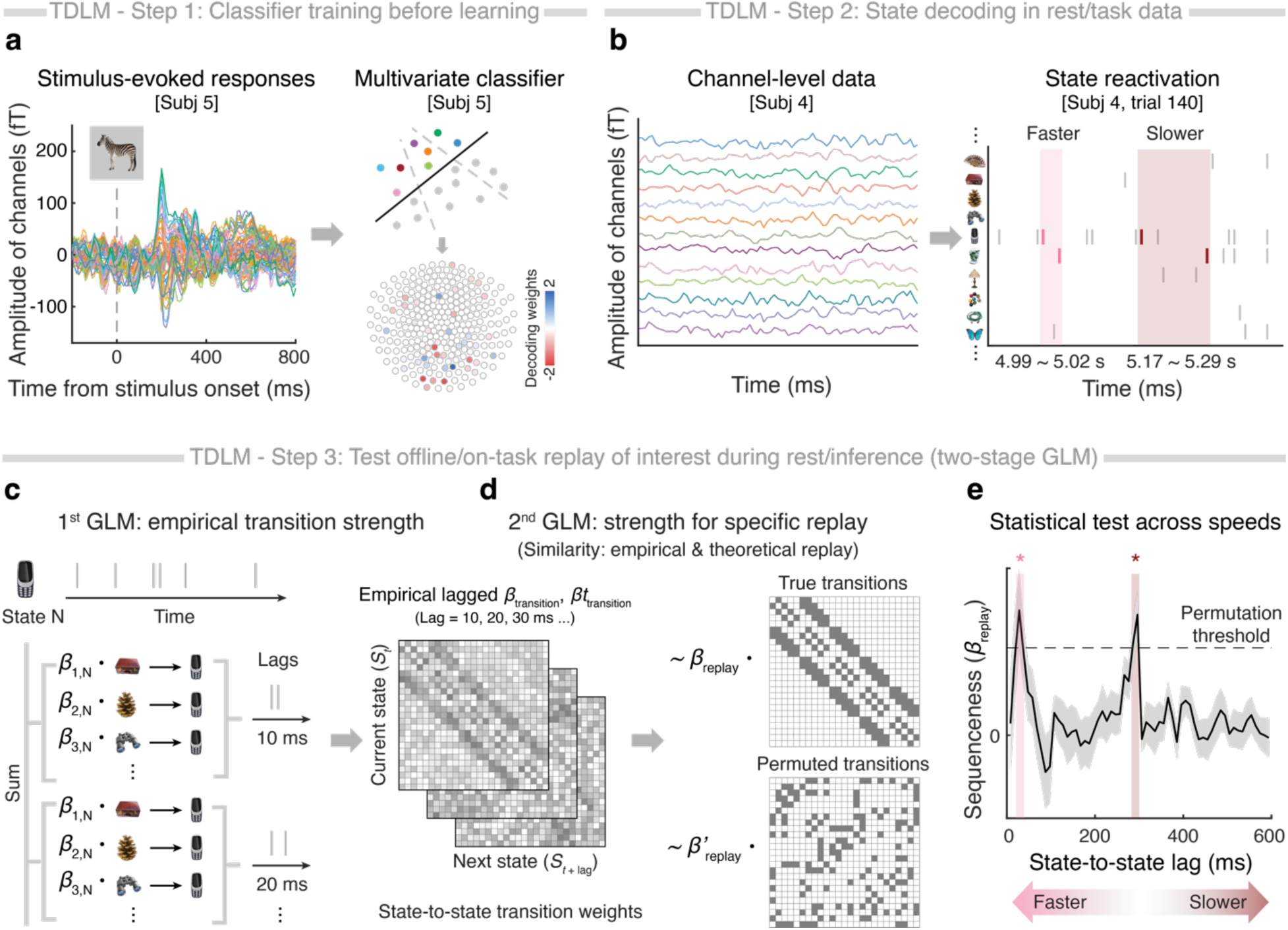
Pipeline for replay detection using temporally delayed linear modelling (TDLM). **a**, During the localizer phase, multi-channel stimulus-evoked data for each stimulus (e.g., zebra, left panel) are used to train binary decoding models via lasso regression, producing channel-specific decoding weights (right panel). **b**, These classifiers are then applied to the channel-level data (left panel) to generate time series of decoded states. The example on the right shows the state reactivation probability for subject 4 during map-based inference. A faster state reactivation (shorter state-to-state lag) is highlighted in pink, while a slower one appears in sangria. **c–e**, Detection of neural sequential reactivations of task-related structures involves a two-stage GLM process. **c**, In the first-stage GLM, we estimate both the empirical strength and its time modulation for each state-to-state transition at various time lags (e.g., 10, 20, 30 ms). **d**, In the second-stage GLM, we derive replay strength (“sequenceness”) and its temporal changes (“sequenceness changed over time”) by mapping these hypothetical transition matrices onto the empirical transition and temporal modulation matrix obtained from the first-stage GLM. **e**, The resulting sequenceness values are plotted against all computed time lags. A non-parametric permutation test (1,000 shuffles) sets the significance threshold across all time lags, controlling for multiple comparisons. Any sequenceness exceeding the 95^th^ percentile of the maximum permuted sequenceness distribution across all time lags is deemed significant. In this simulated example, both fast (pink) and slow (sangria) replay exceed the threshold at distinct time lags.

### Effective neural decoding of all task states

To accurately detect replay using TDLM, we first ensured that neural activations of all task states (29 unique objects) could be reliably decoded. Consistent decodability across states was crucial for unbiased replay detection. Following established protocols (Liu, Dolan, et al., 2021; Liu et al., 2019; Liu, Mattar, et al., 2021; Nour et al., 2021), we trained 29 binary classifiers on task stimuli from the localizer phase using lasso logistic regression (see Methods for details). The highest decoding accuracy occurred when classifiers were trained and tested at the same time point (Fig. 3a, see also Extended Data Fig. 2), peaking at 200 ms post-stimulus onset [10-fold cross-validated accuracy: 44.5 ± 2.6%; chance level: 3.5%].

**Fig. 3.**
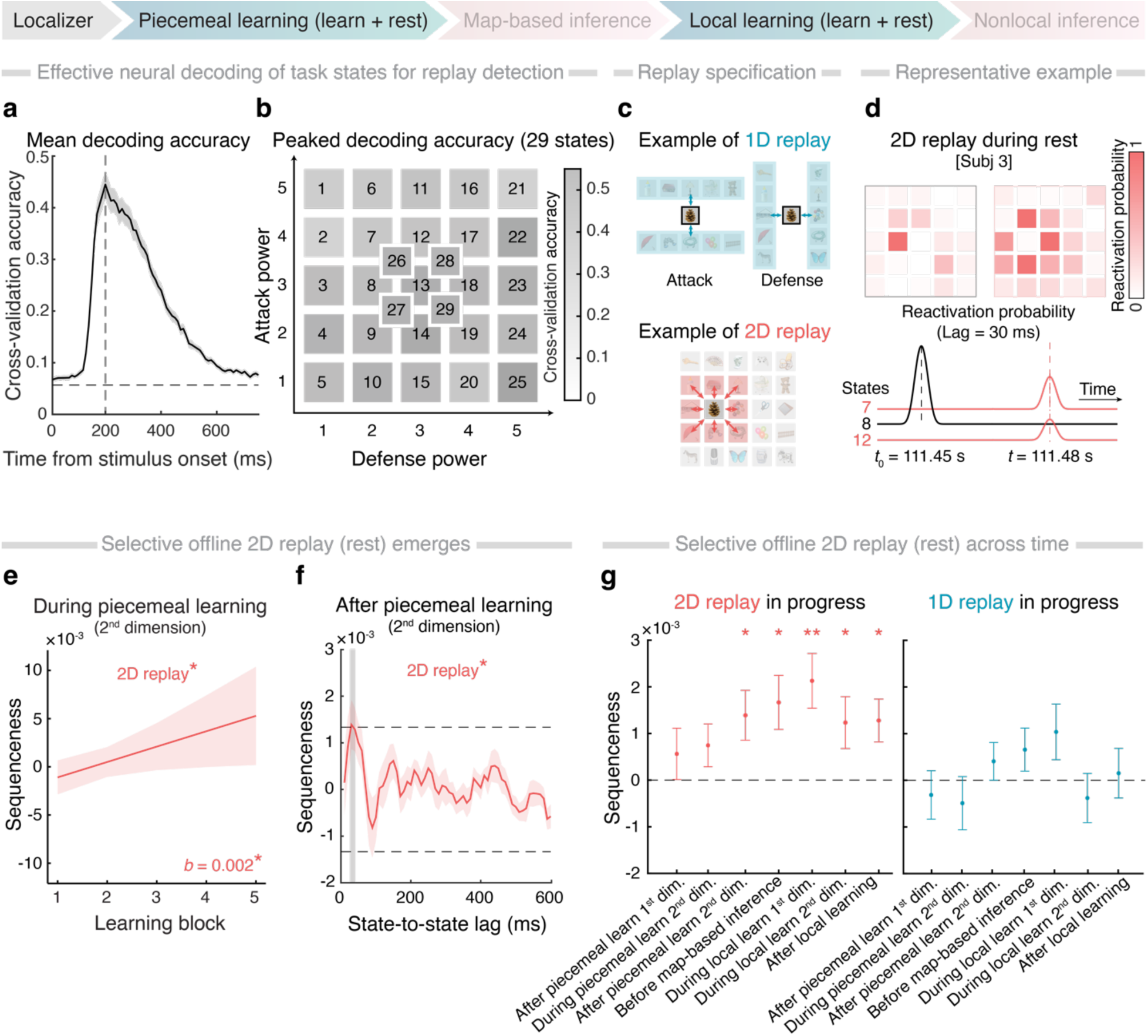
Selective offline 2D replay during and after learning. **a**, Peak mean decoding accuracy across subjects was observed when classifiers were trained and tested at 200 ms after stimulus onset. **b**, No significant difference was found in decoding accuracy across 29 states (numbered by their positions and color-coded by their peak decoding accuracy in **(a)**, see also Extended Data Fig. 2). **c**, 1D replay was defined as neural sequential activations of task states along 1D hierarchies (e.g., attack or defense power), while 2D replay was defined as diffusive transitions from the current state (e.g., “pinecone”) to its surrounding states on the 2D map at a specific time lag. **d**, A representative example of 2D replay was observed during offline rest before map-based inference in subject 3. Empirical reactivation of state 8 (*t*_0_) is followed by activations of adjacent states (e.g., state 7 and 12) in the 2D space after a 30-ms lag. **e**, 2D replay strength (i.e., sequenceness) at a 30-ms time lag increased over the course of piecemeal learning of the 2^nd^ dimension. **f**, 2D replay was significant during post-piecemeal-learning rest. **g**, Summary the offline 2D and 1D replay strength (at a 30-ms time lag) across the experiment, with x-axis labels aligned to the task stages illustrated in Extended Data Fig. 1. The 2D replay was evident immediately after piecemeal learning of the 2^nd^ dimension and remained significant in all subsequent rest periods (left panel, false discovery rate corrected, *ps <* 0.05). However, no significant 1D replay was detected in these same periods (right panel; see also Extended Data Fig. 4). For all figures, the shaded area of the fitted lines in the correlation/regression plots depicts the 95% confidence interval (CI). All error bars and shaded areas represent the standard error of the mean (*S.E.*). Dashed lines represent the non-parametric permutation thresholds. The shaded time windows in the replay graphs highlight the time lags surviving a permutation test. Task phases for the analyses were noted in the experiment schedule at the top. \**p <* 0.05, \*\**p <* 0.01.

Subsequent replay analyses were therefore focused on the multivariate neural representations at this 200 ms post-stimulus onset. Importantly, there were no significant difference in decoding performance across states [repeated-measure ANOVA: *F*(28, 952) = 1.351, *p =* 0.106, Fig. 3b]. Neither attack nor defense power contributed to decoding performance [linear mixed-effect model, fixed effect of attack power: *b* = 0.002 ± 0.006, *p =* 0.753; fixed effect of defense power: *b* = 0.007 ± 0.006, *p =* 0.271] (see details in Methods).

To evaluate the generalizability of decoding across the entire task, we applied these classifiers – trained during the localizer phase – to the subsequent learning and inference phases, and observed robust predicted probabilities [out-of-sample predicted accuracies > 25%; see Extended Data Fig. 3]. These results confirm reliable and comparable decoding performance for replay detection across all task states and time points.

### Selective offline 2D replay accompanies and follows learning

Using our decoding classifiers, we next examined neural reactivation sequences during rest periods interspersed within piecemeal learning. Participants learned pairwise rank relationships separately for each feature dimension (attack or defense). If replay merely consolidated memories, we would expect to observe “1D replay” – sequential reactivation of objects forming linear hierarchies within each dimension. Conversely, if replay supported the integration of information into a 2D map, we would anticipate “2D replay,” where an object is sequentially reactivated alongside its neighboring states on the 2D map at a specific lag (Fig. 3c; see Fig. 3d for a representative example; also Extended Data Fig. 4i). We specifically searched for such 2D replay patterns during the learning of the 2^nd^ dimension, when integration across dimensions became possible.

As participants learned the 2^nd^ feature dimension, spontaneous replay during intermittent rest could, in principle, reflect either within-dimension sequences (1D replay) or integrative cross-dimensional sequences (2D replay). To test this, we modeled both types of replay simultaneously using TDLM. Replay templates included all one-rank transitions along each 1D hierarchy as well as all direct neighbor transitions in the 2D map (Fig. 3c; see Fig. 3d and Extended Data Fig. 4i).

We quantified empirical state-to-state transition strengths and their evolution over time (Extended Data Fig. 7c; see Methods). We found that 2D replay at a 30-ms lag increased across the course of piecemeal learning of the 2^nd^ dimension [linear mixed-effect model, fixed effect of learning block: *b* = 0.0016 ± 0.0007, *p =* 0.035, Fig. 3e; also see Extended Data Fig. 7d]. Notably, 2D replay was already evident immediately after learning the 2^nd^ dimension (Fig. 3f) and remained consistently present in all subsequent rest periods (Fig. 3g, left panel, all *ps <* 0.05, false discovery rate corrected). In contrast, there was no significant evidence of 1D replay at any point during the experiment (Fig. 3g, right panel; see also Extended Data Fig. 4). These replay dynamics cannot be attributed to changes in neural decoding accuracy over time. Together, these findings indicate that offline 2D replay emerges during learning and supports the formation of an integrated task space representation.

### Prioritized offline replay complements learning and supports subsequent inference

Building on the observation of selective offline 2D replay, we explored whether specific parts of the map were prioritized during replay to facilitate representation learning. In the physical space, animals typically explore the boundaries of a new environment first to establish boundary representations (e.g., boundary cells (O’Keefe & Burgess, 1996; Solstad et al., 2008)), before navigating the central areas. We examined whether a similar boundary effect existed in the abstract task space in humans. In our study, boundary objects were defined as those located at the map’s borders, while central objects were situated inside the map border in the 2D space.

Consistent with our expectations, learning performance was significantly better for pairwise relationships involving boundary objects compared to those involving central objects [linear mixed-effect model, interaction of learning dimension (1^st^ vs. 2^nd^) × object position (central vs. boundary): χ^2^(2) = 11.169, *p =* 0.004]. This advantage was found across all learning blocks in piecemeal learning of the 2^nd^ dimension [*b* = -0.392 ± 0.121, *p =* 0.001, Fig. 4a, left]. It also remained significant under a bootstrapped sampling method that accounted for the unequal number of boundary versus central pairs, 95% confidence interval (CI) of mean difference: [0.011, 0.089]. In contrast, no significant difference emerged between boundary and central objects during piecemeal learning of the 1^st^ dimension [*b* = -0.146 ± 0.128, *p =* 0.252]. Moreover, this boundary effect was also significant in the memory test after piecemeal learning (bootstrapped 95% CI of mean difference: [0.005, 0.052]). For visualization, we computed the mean learning performance for each state across participants and then normalized these values to a 0–1 scale in piecemeal learning of the 2^nd^ dimension (see Fig. 4b and Methods). Our behavioral results demonstrate that participants learn the 2D map more effectively at the boundaries than in the central area, similar to patterns seen in spatial navigation.

**Fig. 4.**
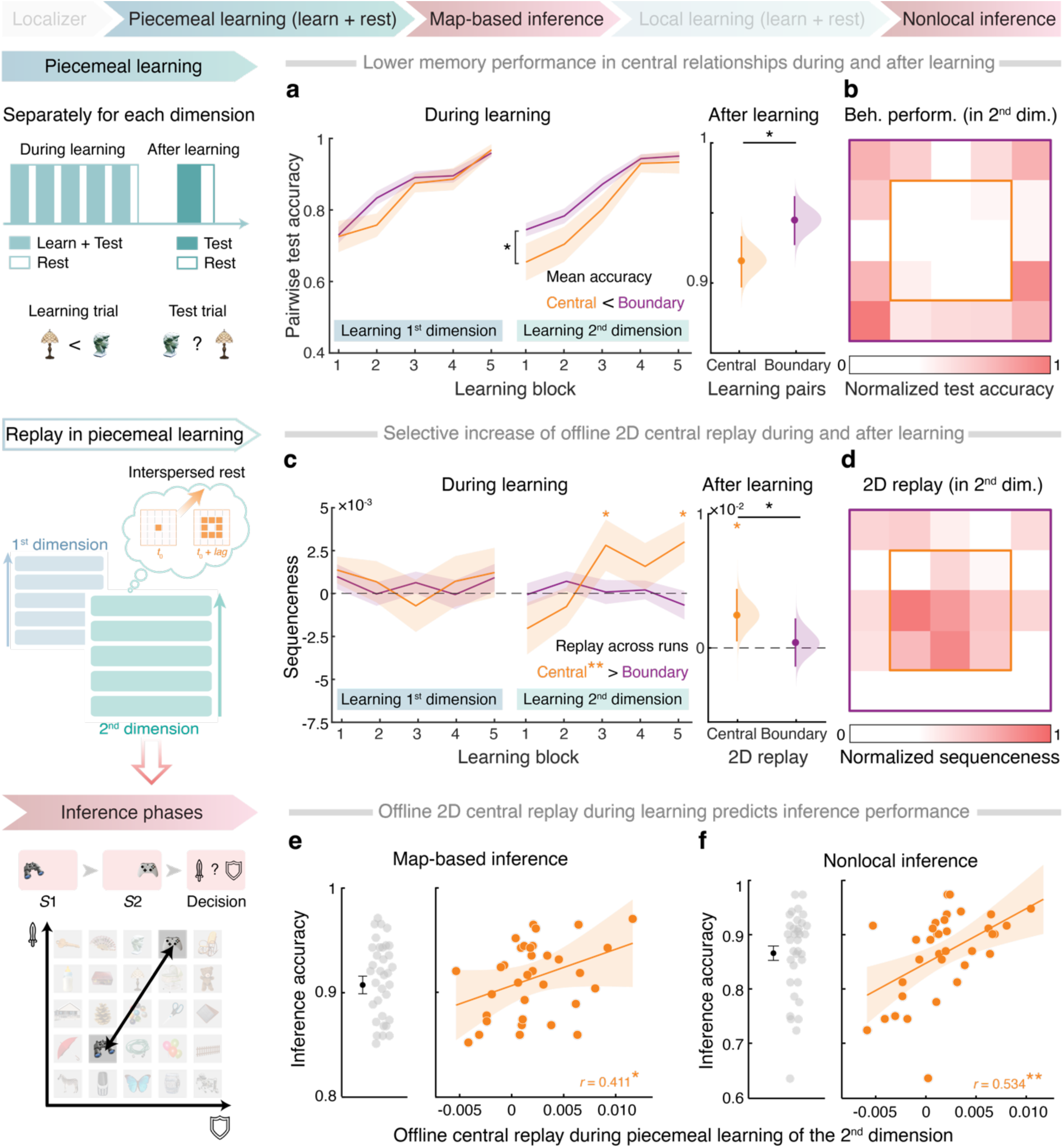
Prioritized offline replay of the central map area complements representation learning and facilitates future inferences. **a**, Higher memory performance for map-boundary (purple) relationships, compared to map-central (orange) ones, was observed during piecemeal learning of the 2^nd^ dimension, but not the 1^st^ dimension (left panel). This difference persisted immediately after learning (right panel, shown with bootstrapped mean distributions and 95% CI). **b**, Normalized memory accuracy in the piecemeal learning of the 2^nd^ dimension was visualized for each task state. **c**, 2D replay of central and boundary relationships across learning blocks revealed no significant increase during the piecemeal learning of 1^st^ dimension. However, a marked increase in central replay at a 30-ms lag, unlike boundary replay, was observed during the 2^nd^ dimension (left panel) and immediately after learning (right panel, see also Extended Data Fig. 5). **d**, Normalized 2D replay strength of task states. **e**–**f**, The strength of offline central replay during piecemeal learning of the 2^nd^ dimension positively correlated with the map-based inference **(e)** and nonlocal inference **(f)** performances. The results remained significant even 1 outlier (3 standard deviations away from the mean, see also Methods) was removed from panel **e** (*r* = 0.357*). \**p <* 0.05, \*\**p <* 0.01.

At the neural level, we examined whether boundary and central relationships differed in 2D replay. Boundary replay was defined as any 2D diffusive transitions involving at least one boundary state, while central replay was defined as transitions between two central states. We modeled both replay types simultaneously in a single GLM, assigning separate regressors for boundary and central transitions, in addition to the 1D replay regressors (see Extended Data Fig. 5a). Contrary to the behavioral pattern, offline replay predominantly represented the central relationships of the 2D map (Fig. 4c–d). Specifically, no significant boundary replay was detected. Instead, central replay increased over time during piecemeal learning of the 2^nd^ dimension [linear mixed-effect model, significant three-way interactions of dimension (1^st^ vs. 2^nd^) × object position (central vs. boundary) × learning block: χ^2^(1) = 4.694, *p =* 0.030; central replay during 2^nd^ dimension: *b* = 0.0012 ± 0.0004, *p =* 0.006; boundary replay during 2^nd^ dimension: *b* = -0.0002 ± 0.0002, *p =* 0.476; Fig. 4c, left]. The central replay was significant during the rest period immediately after piecemeal learning, and was more pronounced than the boundary replay (bootstrapped 95% CI of mean difference: [0.0001, 0.0033], Fig. 4c, right; see also Fig. 4d for normalized 2D replay involving each task state, and Extended Data Fig. 5). This inverse relationship between neural replay and behavioral performance (see also Extended Data Fig. 6) suggests that offline replay prioritizes the weakly-learned information, thereby complementing online learning of the task space (Schapiro et al., 2018).

Further, we assessed whether efficient offline representation learning via central replay contributed to subsequent novel inferences. We found that the strength of 2D central replay during piecemeal learning of the 2^nd^ dimension positively predicted performance of later inference performances, including both map-based inference [Fig. 4e, *r* = 0.411, *p =* 0.029, family-wise error (FWE) corrected] and nonlocal inference [Fig. 4f, *r* = 0.534, *p =* 0.002, FWE corrected].

To validate our findings, we replicated the study using the 2^nd^ task series involving local learning and nonlocal inference. With four new objects introduced to the existing map (see Extended Data Fig. 7b for more details), replay of inferred nonlocal relationships increased preferentially during local learning of the 2^nd^ dimension [nonlocal replay: *b* = 0.0008 ± 0.0003, *p =* 0.017, Extended Data Fig. 7e]. This nonlocal replay predicted future inference performance [*r* = 0.378, *p =* 0.028, Extended Data Fig. 7f, right] even after controlling for memory performance in local learning. Consequently, across both learning phases, offline replay of new knowledge complements online learning in forming effective task representations, facilitating future inference behavior.

#### Distinct on-task replays underlie online planning

Having established the task representation, participants performed novel inferences about the unlearned relationships between objects, either within the existing map (“map-based inference”) or involving newly introduced objects (“nonlocal inference”). On-task replay has been proposed to reactivate experiences or create new links to plan for decision-making (Schwartenbeck et al., 2023; Wimmer et al., 2023). Here we explored the representational content of on-task replays during inference phases. Specifically, in each trial, we examined 2D diffusive replay associated with the first and second stimuli (*S*1-related or *S*2-related transitions), and the trial-specific trajectory of *S*1 – *S*2 (Fig. 5a, d). Both the *S*1-related and *S*2-related (including *S*1 – *S*2) transitions were termed as trial-relevant replay. Trial-irrelevant pairs were defined as all 2D transitions but excluding those related to *S*1 and *S*2.

**Fig. 5.**
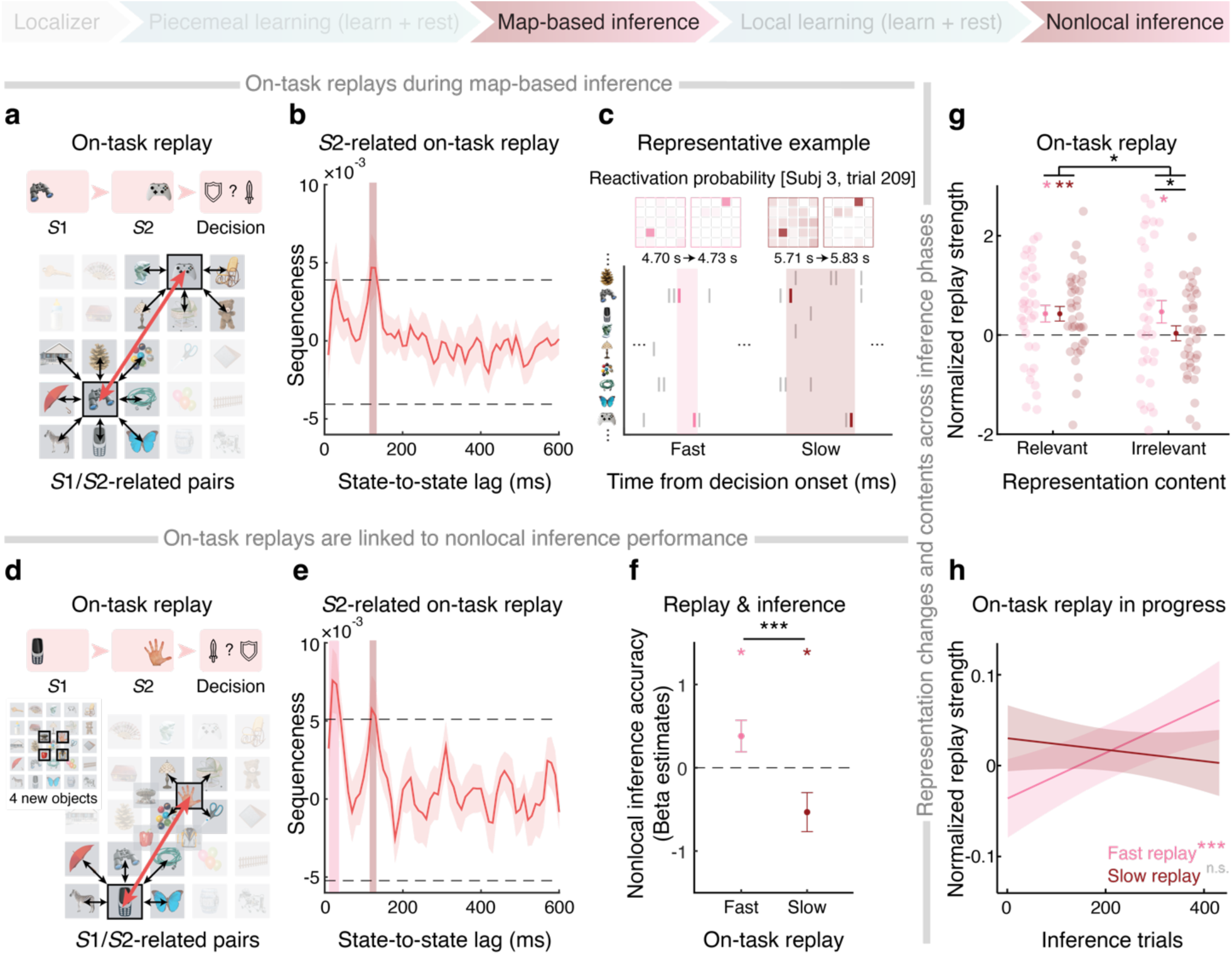
Two different on-task replays underlying novel inferences. **a**, During map-based inference, on-task replay for task-relevant transitions (*S*1- and *S*2-related transitions) is represented by black arrows, and the trial-specific trajectory (*S*1 – *S*2) is depicted in red. **b**, There was significant replay of the *S*2-related transitions, including the trial-specific trajectory, at both a slow speed (120-ms time lag) and a fast speed (30-ms time lag). **c**, Empirical decoded time series with fast and slow replay for the trial-specific trajectory during map-based inference in subject 3 were shown. The normalized reactivation probability of each state at the replay onset and after a specific time lag was highlighted. **d**, Illustration of on-task replay during nonlocal inference. **e**, Similarly, significant 30-ms lagged fast replay and 120-ms lagged slow replay of the *S*2-related transitions were also found in nonlocal inference. **f**, On a trial-by-trial basis, fast on-task replay of the inferred trajectory positively predicted nonlocal inference performance, while slow replay negatively predicted it. **g**, For representational content of on-task replays across inference phases, fast replay represented both trial-relevant and trial-irrelevant pairs (see also Extended Data Fig. 8), while slow replay preferentially represented trial-relevant information. **h**, Across both inference tasks, the strength of fast replay significantly increased with inference trials, which was not true for slow replay. \**p <* 0.05, \*\**p <* 0.01, \*\*\**p <* 0.001.

During map-based inference, we observed significant *S*1-related and *S*2-related replay at a fast speed (peaking at a 30-ms lag, see Extended Data Fig. 8). However, only *S*2-related pairs (including *S*1 – *S*2) showed significant replay at a slower speed (peaked lag of 120 ms, Fig. 5b, see Methods for details). A representative example of fast and slow on-task replays for the inferred trajectory (*S*1 – *S*2) was shown in Fig. 5c.

We observed a similar pattern during nonlocal inference (Fig. 5e; see also Extended Data Fig. 8). In this phase, newly learned objects have not been linked to existing non-adjacent objects previously (Fig. 5d, see also Extended Data Fig. 7b). On a trial-by-trial basis, we found that fast on-task replay for the inferred trajectory *S*1 – *S*2 positively predicted inference performance [linear mixed-effect model, *b* = 0.380 ± 0.189, *p =* 0.045, Fig. 5f], whereas slow replay of *S*1 – *S*2 negatively predicted accuracy [*b* = -0.533 ± 0.234, *p =* 0.023, Fig. 5f], even after controlling for 2D diffusive replays of *S*1- and *S*2-related transitions. These findings suggest the coexistence of two distinct on-task replay processes with opposing impacts on inference performance.

We further examined the representational content of the two on-task replays across both inference phases. Fast replay covered a broader range of 2D diffusive transitions (Extended Data Fig. 8), while slow replay selectively represented trial-related transitions, as evidenced by a significant interaction between replay types (fast vs. slow) and representation content (relevant vs. irrelevant) [repeated-measure ANOVA: *F*(1,34) = 5.346, *p =* 0.027; Fig. 5g]. For trial-irrelevant pairs, only fast replay was significant [fast replay: 0.468 ± 0.227, one-sample *t*-test: *t*(34) = 2.067, *p =* 0.046], and was stronger than slow replay [*F*(1,34) = 4.533, *p =* 0.041]. For trial-relevant pairs, both fast and slow replays were significant [fast replay: 0.429 ± 0.167, one-sample *t*-test: *t*(34) = 2.561, *p =* 0.015; slow replay: 0.427 ± 0.147, one-sample *t*-test: *t*(34) = 2.901, *p =* 0.007], and exhibited no significant difference [*F*(1,34) = 0.0001, *p =* 0.994, Fig. 5g]. This pattern indicates that slow replay primarily concerns trial-specific information, whereas fast replay provides a more general representation of the task.

Intriguingly, we found that fast on-task replay emerged later in each inference trial and its strength gradually increased over time [*b* = 0.0003 ± 0.0001, *p <* 0.001], while slow replay showed no significant temporal changes [*b* = -0.0001 ± 0.0001, *p =* 0.348]. The difference in these temporal slopes was also significant [paired-sample *t-*test: *b* = 0.00002 ± 0.00001, *p =* 0.004, Fig. 5h]. These findings highlight distinct roles of fast and slow on-task replays during novel inference: slow replay focuses on trial-specific computation, whereas fast replay increasingly supports broader task representations over time for efficient inferences.

### Linking offline and on-task replays to grid-cell-like representations of cognitive map

Our findings show that offline replay during rest fosters the formation of an integrated 2D task representation, while on-task replay supports novel inferences: fast replay for general map-level representations and slow replay for trial-specific computations. We next investigated the neural code underlying these representations, focusing on grid-cell-like coding in the EC and mPFC. These regions are known to represent 2D schemas that generalize across environments (Aronov et al., 2017; Hafting et al., 2005). If participants formed such a schema representation in the 2D conceptual space, we would expect to see a grid-like code during inference time, and potentially interacting with different replay types found in our task.

Grid cells fire at regular intervals, forming hexagonal grid fields across space (Hafting et al., 2005; Jacobs et al., 2013). This six-fold periodic firing pattern can be detected using representational similarity analysis (RSA) in human neuroimaging (Bellmund et al., 2016; Constantinescu et al., 2016; Doeller et al., 2010; Park et al., 2021a). Specifically, during mental navigation, we identify pairs of trajectories whose angles differ by roughly 30° (within 15°–45°) as having high dissimilarity in neural activities, capturing the hexagonal grid structure, the six-fold periodicity (Bellmund et al., 2016). First, we computed the neural dissimilarities (1 – *r*) for each pair of trajectories (first-stage GLM), and then mapped these empirical dissimilarities onto the theoretical matrix encoding six-fold periodicity (second-stage GLM). The resulting beta weights tracked the strength of grid-like coding over the mental navigation time course.

We found a significant grid-like code in source-localized EC and mPFC after *S*2 onset [390–980 ms, *p =* 0.030, cluster-level FWE corrected, Fig. 6b, left]. This effect specifically aligned with six-fold symmetry (*b* = 0.0018 ± 0.0007, *p =* 0.044, FWE-corrected; Fig. 6b, right), with no other periodic patterns observed in control regions (e.g., primary visual/motor cortices) or at different frequencies (Extended Data Fig. 9). We also tested four-, five-, seven-, and eight-fold symmetries and found no significant effects (Extended Data Fig. 9), confirming the specificity of the hexagonal pattern. Moreover, this grid-like schema generalized to a “new” map during nonlocal inference (Extended Data Fig. 10), indicating a stable code for the 2D conceptual space.

**Fig. 6.**
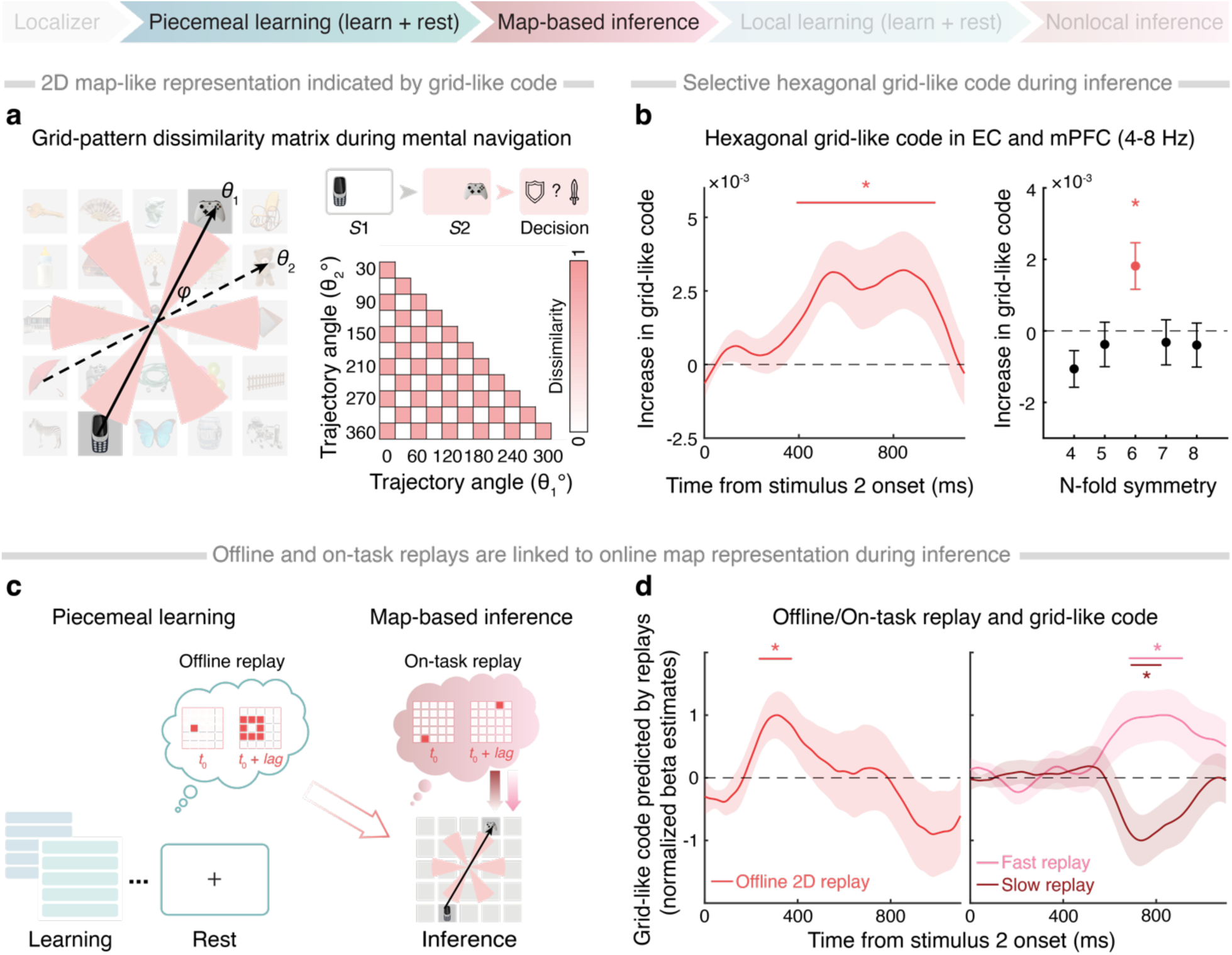
Neural representation of the cognitive map and replays. **a**, During mental navigation, inferred trajectories traverse the 2D map and intersect hexagonally arranged grid-cell firing fields. Because of this six-fold symmetry, trajectories at approximately 30° apart (e.g., *θ*_1_ and *θ*_2_) exhibit more dissimilar neural activity than those at 0° apart. Following Bellmund et al. (2016), we used an RSA-like approach to detect the hexagonal pattern based on these angle differences (*φ*). **b**, A selective hexagonal grid-like code was observed in the entorhinal and medial prefrontal cortices after *S*2 onset (390 to 980 ms) in the theta-band (4-8 Hz) of neural activity (see also Extended Data Fig. 9). Such grid-like code had specifically six-fold symmetry, after averaging over the *S*2 presentation time (right panel). **c**, Schematic illustration of the role of offline replay before map-based inference and on-task replay during map-based inference in grid-like representation. **d**, For offline replay during rest, the grid-like representation was positively correlated with the 2D replay (left panel). For on-task replay, the grid-like code was positively associated with the fast replay, but negatively correlated with the slow replay (right panel). The line with asterisks above represents the time windows (cluster-level FWE-corrected, *p <* 0.05). \**p <* 0.05.

Having identified a grid-like representation of the 2D space, we then investigated its link to different types of replays, both offline and online (Fig. 6c). Specifically, we tested whether offline replay before map-based inference helped build the grid-like code and whether the grid-like representation facilitated inferences about unlearned relationships. We found that this offline replay positively predicted the emergence of grid-like code online shortly after *S*2 onset [significant time window: 230-380 ms, cluster-level FWE corrected, Fig. 6d, left], suggesting that offline replay facilitated construction of the grid-like schema representation. Later in each trial, fast on-task replay showed a positive relationship with the grid-like code [680–920 ms], whereas slow on-task replay showed a negative relationship [690–820 ms, cluster-level FWE corrected; Fig. 6d, right]. Considering that fast replay encodes the general 2D map while slow replay represents only the trial-relevant information, the distinct correlation patterns between on-task replays and grid-like code implied that a well-established schema representation of the task space can reduce the need for slow, task-specific computation.

Besides, we conducted multiple control analyses to rule out potential confounds, such as changes in head positions or orientations. No significant correlation was found between the relative head positions/orientations and task-related neural measures such as offline replay, on-task replay, or cognitive map representations (Extended Data Fig. 11).

Together, these findings link different types of replays to the grid-cell-like code of the cognitive map, highlighting their distinct roles in structuring the 2D map from piecemeal experiences and using the map for flexible inference.

### Neural signatures underlying offline and on-task replays

In light of the distinct functional roles of replay in offline representation learning and online inference, we characterized the neurophysiological signatures of both offline and on-task replays. In both rodents and humans, offline replay events often coincide hippocampal sharp-wave ripples (Buzsáki, 2015; Liu et al., 2019). Consistent with previous findings, in the learning phases, we found a significant power increase in the ripple frequency band of 120-150 Hz at the offline replay onset during rest, compared to the pre-replay baseline (Fig. 7a). This increased power was linked to enhanced source-localized activity in the medial temporal lobe (MTL), including the hippocampus [cluster-level FWE corrected, *p <* 0.05, peak coordinates: *x =* -21, *y* = -1, *z* = -46; Fig. 7b]. For on-task replay during inference, there was a significant increase in the ripple power at the onset of fast replay [0.012 ± 0.003, one-sample *t*-test: *t*(34) = 4.089, *p <* 0.001], but not of slow replay [0.005 ± 0.003, one-sample *t*-test: *t*(34) = 1.494, *p =* 0.144]. Such difference was also significant in both parametric and non-parametric tests [paired-sample *t*-test: *t*(34) = 2.052, *p =* 0.048; Wilcoxon signed-rank test (more robust to outlier): *z* = 2.162, *p =* 0.030, Fig. 7c].

**Fig. 7.**
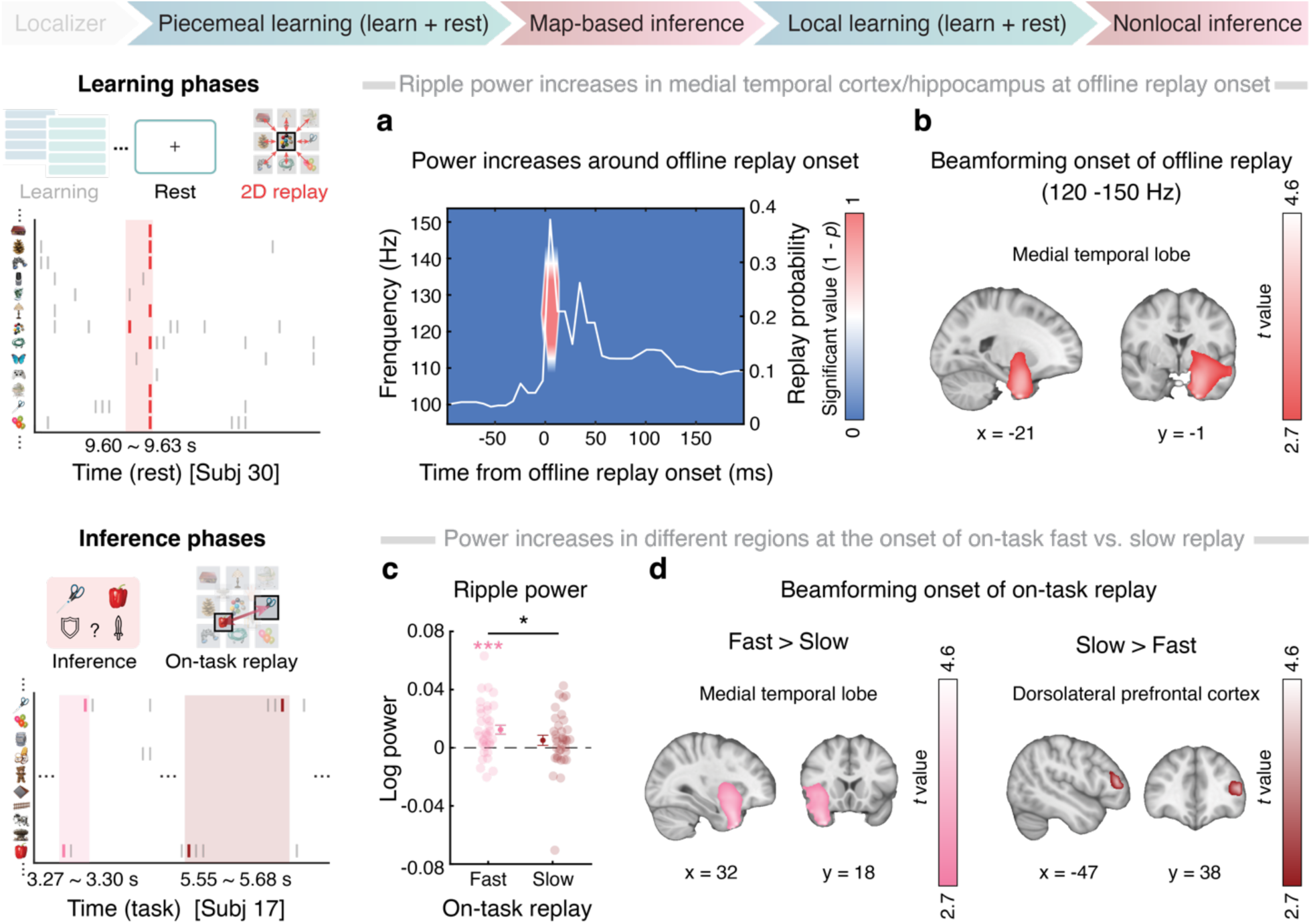
Neural signatures of offline and on-task replays. **a**, Across resting periods after piecemeal learning (see empirical examples on the left), there was a significant increase in ripple power (120-150 Hz) at offline replay onset (cluster-level FWE-corrected *p <* 0.05), aligning with the peak of replay probability (white line). **b**, Source localization of ripple frequency band power at the replay onset showed significant activations in the anterior temporal lobe, including the hippocampus (peak MNI coordinates: x = -21, y = -1, z = -46). **c**, Similar to the offline replay in A, the fast on-task replay was also linked to ripple power increase at replay onset during inference phases. This power increase was significantly stronger than that observed in the slow replay, which, on its own, was not significant. **d**, For source localization of gamma band power (80-180 Hz) at the replay onset, significantly stronger activation in the medial temporal lobe, including the hippocampus, was found for fast on-task replay, compared to slow replay (peak MNI coordinates: x = 32, y = 18, z = -41, cluster-level FWE-corrected, *p <* 0.05). Stronger activations in the dorsolateral prefrontal cortex were observed for slow on-task replay, relative to fast replay (peak MNI coordinates: x = -47, y = 38, z = 12, cluster-level FWE-corrected, *p <* 0.05). \**p <* 0.05, \*\*\**p <* 0.001, *n.s*.: non-significant.

To ensure an unbiased comparison of neurophysiological features between fast and slow on-task replay, we expanded our analysis to a broader ripple-band range (80–180 Hz), often referred to as “high gamma”, following previous work (Liu, Mattar, et al., 2021; Veselic et al., 2025). This approach avoids restricting the analysis to the narrower ripple frequencies typically linked with fast replay. Fast replay was associated with a localized increase in gamma power in the MTL [cluster-level FWE corrected, *p <* 0.05, peak coordinates: *x =* 32, *y* = 18, *z* = -41; Fig. 7d, left]. Conversely, slow replay showed higher gamma power in the dorsolateral prefrontal cortex (dlPFC) than fast replay [cluster-level FWE corrected, *p <* 0.05, peak coordinates: *x =* -47, *y* = 38, *z* = 12; Fig. 7d, right]. These results suggest that offline replay and fast on-task replay share similar neurophysiological features, potentially supporting general representations of the 2D map. In contrast, slow on-task replay appears to be selectively involved in deliberate, task-specific computations.

## Discussion

This study provided unprecedented insights into how the human brain constructed a 2D cognitive map from fragmented memories and then used this map for flexible inferences. During resting periods interspersed throughout the learning process, offline replay of the inferred 2D task structure – rather than simple 1D memories – became increasingly prominent, prioritizing weakly learned relationships. This offline replay, in turn, facilitated the formation of grid-cell-like map representations and supported subsequent novel inferences. In the inference phase, the grid-like code served as a schema across two different maps, reducing the need for slower, task-specific computations carried out by on-task replay.

Our findings offer convergent evidence for offline replay in humans, highlighting both its rapid speed (∼30-ms state-to-state transitions) and its hippocampal involvement, marked by ripple-band signatures in nonspatial cognitive tasks. In both learning phases, we observed spontaneous 2D replay emerging during rest with this compressed timescale, accompanied by increased ripple-band power in the source-localized MTL, including the hippocampus. These findings align with recent single-neuron recordings in humans, where hippocampal neurons replay inferred relational structures at a ∼30-ms timescale (Tacikowski et al., 2024). As hippocampal replay is characterized by sharp-wave ripple events in rodents (Buzsáki, 2015; Roux et al., 2017), we also observed increased ripple-band power around replay onset, replicating earlier reports of human replay in linear task spaces (Liu et al., 2019; Liu, Mattar, et al., 2021; Nour et al., 2021). Our recent iEEG study further demonstrated that human hippocampal ripples during rest correlated with the learning of a 2D conceptual map and predicted its grid code in the EC (Xiao et al., 2025). These results suggest that the offline replay measured by MEG shares both anatomical specificity and functional characteristics with hippocampal replay across species.

Although extensive studies have highlighted the roles of hippocampal place cells and entorhinal grid cells in representing cognitive maps (Moser et al., 2008), few have probed the underlying learning processes that form such maps. We found that offline replay reorganized piecemeal memories into a unified map-like structure, thus linking replay to the formation of a 2D cognitive map. Notably, even when participants were exposed only to 1D information without explicit awareness of the 2D task space, replay extended beyond direct memories to infer 2D configurations during rest. A similar process emerged in local learning when new objects were introduced to an existing map: increased nonlocal replay between these new objects and non-adjacent existing ones predicted performance in future novel inferences. Consequently, offline replay infers new relationships from learned fragmented experiences, building a more complete cognitive map that supports flexible behavior.

Intriguingly, offline replay also complements online learning by targeting weakly learned information rather than representing all states uniformly. During piecemeal learning, replay predominantly involves the central area of the 2D map, an adaptive approach that strengthens relationships with lower performance, thereby fostering a fully integrated 2D map. Boundary pairs, being simpler and more salient, tend to reach near-ceiling performance early on, leaving little room for replay-driven improvement. By contrast, central objects demand more complex integration, so replay focusing on these less stable regions yields greater benefits. Recent animal research indicated that replay similarly overrepresented less-experienced trajectories during rest and sleep (Gillespie et al., 2021), and one explanation is that replay protects the rich internal task representation from unbalanced online experiences (H. T. Chen & Van Der Meer, 2024; Courellis et al., 2024). Although those studies focused on replays after task representations had been largely established, the idea aligns with our findings: offline replay supplements online experiences to build balanced task representations. It further suggests that new place fields might form after offline replay when we are learning a new task space (Bakermans et al., 2025).

Our observation of 2D “diffusive” replay differs from the more linear, path-like replay often reported in animals navigating physical spaces. Rather than focusing on a particular path, these replays spread in multiple directions across the 2D conceptual map. It is possible that our 2D learning procedure, unlike physical navigation, produces replay patterns spanning a broader range of transitions instead of adhering to single, linear trajectories. Indeed, the data show that replays do not appear restricted to a specific direction; rather, they involve various connections in the conceptual map. This aligns with Brownian diffusion-like sequences thought to be optimal for learning new spaces (McNamee et al., 2021; Stella et al., 2019). Future studies with higher temporal resolution or complementary recording techniques (e.g., iEEG or single-unit data) could clarify whether these diffuse patterns arise from multiple distinct replay events in parallel or if certain transitions predominate under specific contexts.

After offline representation learning, the resulting cognitive map was characterized by a hexagonal, grid-like code in the source-localized EC and mPFC during inference. Acting as a schema, this code generalized across two maps and supported novel inferences, paralleling earlier results from human spatial and mental navigation (Doeller et al., 2010; Horner et al., 2016; Jacobs et al., 2013; Jacobs & Lee, 2016; Xiao et al., 2025). We also observed that offline replay appeared to initiate early neural processes of grid-like representation, suggesting that it might “preconfigure” the 2D map structure for efficient inferences.

In both inference phases, we identified two types of on-task replay – fast and slow. Fast replay, occurring at a ∼30-ms state-to-state lag, represented the entire 2D map (including both task-relevant and irrelevant transitions) and correlated positively with inference performance as well as the grid-like code. It parallels hippocampal replay observed across species, exhibiting a similar speed and neural characteristics to the offline replay identified during rest in this study and in previous MEG work (Liu et al., 2019; Liu, Mattar, et al., 2021), as well as recent invasive neuronal recordings in the human MTL (Tacikowski et al., 2024; Xiao et al., 2025). Notably, fast replay emerged later in the inference process, presumably reflecting an increasing reliance on the internal map representation to infer unobserved relationships. As participants become more familiar with the map, they navigate it more efficiently, thereby improving inference performance. Consistent with this, hippocampal ripples observed in a human iEEG study have also been shown to predict later emergence of entorhinal grid-like code and inference performance (Xiao et al., 2025).

By contrast, slow on-task replay, at a ∼120-ms state-to-state lag, focuses on trial-specific information and appears throughout the inference phase – even in earlier stages when participants are still learning to apply the cognitive map. This slower replay is not associated with hippocampal ripples but is source-localized to the dlPFC. Rodent and primate studies similarly suggest that PFC neurons encode task-relevant specifics, such as relative positions from start to goal (El-Gaby et al., 2024), and that frontal replay can be independent of hippocampal replay, yet varies with cognitive demands (Kaefer et al., 2020). In our data, slow replay strength is negatively related to the grid-like code, indicating it may serve as a fallback mechanism for more deliberate, trial-by-trial computations, akin to working memory maintenance in frontal regions (Bastos et al., 2018; Lundqvist et al., 2018). These complementary replay modes thus allow the brain to shift from slower, detail-focused processing to faster, more “intuitive” inference as learning and familiarity advances.

Finally, several limitations should be acknowledged. Although we detected increased ripple-band power at replay onset, MEG is inherently less sensitive to high-frequency oscillations, particularly from deep-brain structures like the hippocampus, owing to the 1/*f* nature of MEG power spectra. While our findings broadly match neurophysiological evidence from human single-neuronal and intracranial local field potential recordings, direct interpretations of hippocampal replay in MEG alone must be approached with caution. In addition, we employed MEG source modeling to explore the neural sources of both offline and on-task replay events. However, the inverse problem in MEG source localization (Hillebrand & Barnes, 2002) complicates the precise identification of deep brain generators, including the EC. Even so, the recent complementary iEEG evidence (Xiao et al., 2025) strongly supports the role of hippocampal ripples in forming and using cognitive maps. A selective theta-band grid-like code has also been confirmed in the EC and mPFC with invasive recordings, reinforcing our MEG-based findings.

In conclusion, our results offer a comprehensive account of how the human brain integrates fragmented experiences to optimize learning, memory, and decision-making in a 2D conceptual space. Offline replay complements online experiences for efficient schema formation, reducing the need for deliberate on-task computations during inference. This dynamic interplay between offline and on-task replays enables the brain to generate new knowledge and flexibly generalize it to future scenarios.

## Methods

### Participants

Forty participants in total were recruited through university advertisements and social media. Five were excluded (four chose not to continue, one provided random responses), resulting in a final sample of 35 healthy adults (22 males, 13 females; mean age 23.94 ± 2.40 years, range 20–31). An a priori power analysis indicated that 34 participants would be needed to detect a significant difference from zero in a two-tailed one-sample *t*-test [effect size *d* = 0.5, *α* = 0.05, power = 0.80]. All participants were right-handed, had normal or corrected-to-normal vision, and reported no history of psychiatric or neurological disorders or metal implants. They provided written informed consent in accordance with the Declaration of Helsinki, under approval by the Institutional Review Board of the State Key Laboratory of Cognitive Neuroscience and Learning at Beijing Normal University (ethics number IRB_A_0051_2022002). Participants were advised to have enough sleep and avoid alcohol and caffeine on the day before the MEG scanning session, which was followed by a separate MRI session approximately one week later for structural imaging. They received compensation of ¥120 per hour plus a performance-based bonus ranging from ¥200 to ¥500.

### Stimuli and task design

The task was designed to explore the gradual formation of a five-by-five mental map and assimilation of new knowledge. Further, we investigated the role of the constructed representations in novel inferences. Initially, 25 objects were assigned to different unique locations on the 2D map along two independent feature dimensions (attack power and defense power). The experiment was divided into in five phases, consisting of two task series of learning and inference phases (see Fig. 1 and Extended Data Fig. 1): (**i**) Localizer: Identifying neural representations of all task states; (**ii**) Piecemeal Learning: Learning pairwise rank relationships between objects within each dimension separately (attack or defense power); (**iii**) Map-based Inference: Inferring unobserved relationships between objects, either within a single dimension (1D inference) or across two dimensions (2D inference); (**iv**) Local Learning: Assimilating new knowledge of local relationships between four newly introduced objects and their neighbors, separately for each learning dimension; (**v**) Nonlocal Inference: Deducing unlearned relationships between newly introduced objects and other nonlocal objects on the 2D map. Each task phase was divided into multiple scanning runs lasting approximately 8 minutes. In both learning phases, 1-mintue resting periods interspersed during learning and 5-minute rest periods after learning phases were used to examine how offline replay contributed to building and maintaining the cognitive map (see Fig. 1 and Extended Data Fig. 1). Approximately one week after the MEG session, participants underwent an MRI scan for source localization.

#### Localizer phase

Similar to previous studies (Liu, Dolan, et al., 2021; Liu et al., 2019; Liu, Mattar, et al., 2021; Nour et al., 2021), this task phase was used to train stimulus classifiers for detecting their sequential neural reactivations during rest and tasks. To prevent exposure to the task structure during classifier training, this phase was performed before learning. To ensure equivalent decoding accuracy in different states, the mapping from pictures to task states was held constant within subjects but randomized between subjects. Pictures were presented in a randomized order in 6 runs, and each picture was shown for 45 trials. Specifically, in each trial, a short fixation period “+” was shown for 0.2 s. Then, a picture (e.g., balloon) occurred at the center of the screen for 0.8 s. After that, a text label (e.g., bear) appeared, and participants were asked to determine the correspondence of the text to the preceding picture within 1 s (see Extended Data Fig. 1, upper panel). This probe was to encourage semantic representation of the picture rather than visual encoding only (Kurth-Nelson et al., 2016).

#### Piecemeal learning phase

During piecemeal learning, participants were tasked with memorizing pairwise relationships between objects with a one-rank difference in each dimension. For each object, there were two feature dimensions: attack power and defense power. The attack and defense power of each object was independent and bore no relation to real-life experiences. In the piecemeal learning phase, two dimensions were learned sequentially. Each displayed pair differed by only one rank in the given dimension. Importantly, participants were not exposed to the true structure of the task space, nor were they instructed to link stimuli to a hierarchy or to a 2D map. This approach was similar to previous training protocols of building cognitive map through piecemeal learning (Park et al., 2020, 2021b, 2021a).

Participants learned pairwise relationships for two separate dimensions in a randomized order. To assess learning progress both online (during tasks) and offline (during rest), each dimension was structured into five runs. Each run comprised a 5-minute “*<u>learn</u>*”, a 2-minute “*<u>test</u>*” and a 1-minute “*<u>rest</u>*” (see Fig. 1, Fig. 4 top-left panel). Note that within each “*<u>learn</u>*” block, subjects were exposed to all 25 objects. The visual experiences of object pairs remained the same in each learning block of each dimension, with each pair appearing twice per dimension. This design ensured that any observed replay differences were not due to variations in experiences (see Extended Data Fig. 1, upper panel, and also Extended Data Fig. 7a).

In the “*<u>learn</u>*” stage, 20 pairs of objects were presented twice in a random order to ensure unbiased learning. Each pair of objects had only one-rank difference in the given dimension. Each trial began with a short fixation period (0.2 s). Next, two objects were presented, one by one, at the center of the screen for 1 s. Between the two objects, a symbol “>” or “<” occurred for 0.8-1 s, indicating the relative rank between two pictures in the given dimension. After a short inter-stimulus interval (0.5-0.7 s), participants were quizzed on the relationship between the two objects shown previously (objects were placed randomly in the left and right side). They were asked to select the one with a stronger power in the given dimension within 2 s. The selected option was framed with a square. Then, feedback “√” or “×” appeared, followed by an intertrial interval of 0.6-0.8 s.

During the “*<u>test</u>*” stage, participants were quizzed on the 20 pairwise relationships without receiving feedback, to avoid further learning at this stage. In each trial, two objects were presented randomly in either side one by one for 1 s with an inter-stimulus interval of 0.8-1 s. Then, a decision rule indicating the currently learned dimension (i.e., a “sword” icon for the attack power or a “shield” icon for the defense power in both sides) appeared at the positions where the objects were presented. Participants were asked to select the stronger one in the given dimension within 3 s.

During the “*<u>rest</u>*” stage, participants were instructed to remain still, keep their eyes open, and focus on a central fixation “+” on the screen for 1 minute. This stage aimed to detect the existence as well as the progress of offline learning of the cognitive map.

After 5 runs of piecemeal learning in each dimension, participants were tested on memories of the 100 pairs learned previously (pairwise test). No feedback was shown. The trial order was randomized. Additionally, participants went through a 5-minute rest session prior to between-session break for ∼20 minutes. Moreover, to test whether participants memorized the relative ranks in both dimensions clearly and recalled them flexibly, we designed an intermixed test in which the learned pairwise relationships in both dimensions were interleaved randomly across trials (see Extended Data Fig. 1, middle left). The intermixed test required the ability to flexibly retrieve memory from each dimension. This test assessed the memory performances on both dimensions. No feedback was provided. The intermixed test was administered three times during the experiment: once after the post-piecemeal-learning rest period, and then once before the relearning phase (after nap) and once after the relearning phase. The relearning phase was developed to avoid memory forgetting before map-based inference (see Extended Data Fig. 1, middle panel). In this phase, participants revisited the relationships they had previously answered incorrectly with a fixed number of repetitions. In each trial, the pair of objects were presented in either side of the screen, with a “>” or “<” symbol and a “sword” or “shield” icon indicating the rank information and learning dimension, respectively.

#### Map-based inference phase

As participants were familiar with the pairwise relationships between objects, we then tested whether participants were able to infer unobserved relationship. We developed a map-based inference task, separately for 1D and 2D inferences (see Extended Data Fig. 1, middle panel). For 1D inference, participants were asked to select the object with a higher attack or defense power (cued by the two icons of “sword” or “shield”, respectively). For 2D inference, participants were required to identify whether the attack power of one object exceeded the defense power of the other based on the decision rule (e.g., “sword” vs. “shield” symbols signified the comparison between the “attack power” of the object presented on the left and the “defense power” of the object on the right). In both tasks, two objects occurred, one by one, in either side of the screen for 1 s, with an inter-stimulus interval of 0.7-0.9 s. The presented order of each pair of objects was randomized. After a short inter-stimulus interval (0.5-0.7 s), a decision rule (i.e., two icons indicating the corresponding dimension for each object) was shown and participants were required to decide within 3 s following the rule. No feedback was provided to avoid further learning.

If participants constructed a 2D mental map, the transition from one object to another is like drawing a trajectory on the map (see Grid-like code for details). To ensure an unbiased estimate of grid-like code of the map (Constantinescu et al., 2016; Park et al., 2021a), the angles of these sampled trajectories were evenly distributed across 24 bins, with each bin spanning 15 degrees. At the end of the map-based inference phase, participants performed a position test on the location of each object on the map (see Extended Data Fig. 1, middle right panel). This was to probe the explicit knowledge of the 2D map. The mean accuracy was 0.756 ± 0.018, with no difference between dimensions [1^st^ dimension: 0.753 ± 0.021, 2^nd^ dimension: 0.759 ± 0.019; paired-sample *t*-test: *t*(34) = 0.344, *p =* 0.733], indicating an overall good knowledge of the 2D map.

#### Local learning phase

Having identified the cognitive map was built, we next introduced four new objects into the central area of the existing map (see Extended Data Fig. 1, bottom-left panel, and also Extended Data Fig. 7b). Specifically, participants were shown 16 pairs of local relationships between new objects and their neighbors on the map, twice for each dimension. The order of learning dimensions was counterbalanced between subjects. Similar to the training procedure in piecemeal learning, during local learning, participants went through a 4-minute “*<u>learn</u>*” phase, a 2-minute “*<u>test</u>*” phase and a 1-minute “*<u>rest</u>*” phase (see Extended Data Fig. 1, bottom-left panel). After 4 blocks of training in local learning, there was also a 5-minute resting period.

#### Nonlocal inference phase

After post-local-learning rest, participants were invited to infer unexperienced relationships between the new object and existing nonlocal ones (not neighbors) on the 2D map. Similar to the map-based inference, participants were required to infer whether the attack power of one object exceeded the defense power of the other one based on a decision rule (see Extended Data Fig. 1, bottom panel). No feedback was provided. In line with those in 2D inference, the angles of the sampled trajectories were evenly distributed across 24 bins, with each bin spanning 15 degrees.

### MEG data acquisition and preprocessing

The MEG data were collected at 600 samples/s using a whole-head 275-channel axial gradiometer system (CTF Omega, VSM MedTech). The task was displayed on a screen through a mirror projection. To ensure high data quality, we designed shorter recording runs, scheduled regular breaks, and maintained consistent head positioning throughout the session.

During each MEG scanning run, participants sat upright and were instructed to remain still and relaxed. Short breaks of one to two minutes were given between runs to minimize fatigue and reduce muscle-related artifacts. In addition, after every hour of scanning, participants took a longer break of approximately 20 minutes if needed.

Prior to scanning, we marked standard anatomical reference points (nasion and left/right preauricular points) and placed head localization coils accordingly. Participants’ head positions were recorded at the start of each session. At the beginning of each run, they were asked to reposition their heads to match the initial location and orientation, verified using the MEG system’s localization procedure. MEG recording was paused if excessive head motion or muscle artifacts were detected, ensuring stable and high-quality data collection.

Following the preprocessing pipeline established in our previous studies (Liu, Dolan, et al., 2021; Liu et al., 2019; Liu, Mattar, et al., 2021; Nour et al., 2021), MEG data from all channels were first high-pass filtered at 0.5 Hz using a bidirectional first-order IIR filter to remove low-frequency drift. The data were then downsampled to 400 Hz and inspected for bad segments caused by excessive noise. To improve artifact detection, data from multiple scanning runs within each task session were concatenated for independent component analysis (ICA) using the FastICA algorithm (http://research.ics.aalto.fi/ica/fastica/). ICA decomposed the data into 150 temporally independent components (default setting in OSL), and their associated sensor topographies. Artifact components were identified through manual inspection of each component’s spatial distribution, time course, and frequency spectrum, and were subsequently removed.

After ICA, we re-inspected the data to identify and exclude any remaining bad channels or segments, resulting in a cleaned MEG dataset. For most subsequent analyses – such as decoding, reactivation, and sequence detection – the cleaned MEG data were further downsampled to 100 Hz to improve signal-to-noise ratio. Exceptions included source reconstruction, time-frequency analysis, and grid-like code analysis, which were conducted using the 400-Hz cleaned data. Unless otherwise noted, all analyses were performed at the whole-brain sensor level, with MEG signals expressed in femtotesla (fT).

### MRI data acquisition

MRI scans were performed on a 3T Siemens MAGNETOM Prisma MRI scanner. Structural data for each individual were acquired using a T1-weighted magnetization-prepared rapid acquisition gradient echo sequence with the following parameters: TR = 2,530 ms; TE = 2.98 ms; TI = 1,100 ms; flip angle = 7°; voxel size = 0.5 mm × 0.5 mm × 1 mm; field of view = 256 mm × 256 mm. The individual T1-images were used to co-register to the MEG coordinates system before source localization (see also Source localization below).

### Behavioral analysis

In the piecemeal learning phase, to measure the learning effects on the pairwise relationships in different dimensions, we used a generalized linear mixed-effect logistic regression to model the trial-by-trial test accuracy (binary variable, 0 or 1) across five runs in both dimensions. In particular, the fixed-effect variables were the learning block, learning dimension, and their interaction with central/boundary pairs as fixed effects.

### Stimulus decoding

Sequenceness analyses during rest and tasks relied on the ability to quantify evidence of transient spontaneous neural reactivations of task stimuli from multivariate patterns of MEG sensors. Only the sensors that were not rejected across all runs were then used for classifier training. Following our established approach (Liu, Dolan, et al., 2021; Liu et al., 2019; Liu, Mattar, et al., 2021), we trained separate lasso-regularized logistic regression models for each of the 29 stimuli with the evoked neural responses during the localizer task (Fig. 2a). The training and testing timepoints ranged from stimulus onset to 750 ms post-stimulus onset. A training scheme using a single time point of a 10-ms bin was employed to mitigate potential temporal confounding in replay detection (Liu, Dolan, et al., 2021; Vidaurre et al., 2019). Inclusion of null data from pre-stimulus onset as negative examples facilitated sequenceness detection by reducing the spatial correlation between classifiers (Liu, Dolan, et al., 2021). The decoded probability was transformed using the nonlinear sigmoid function. We used L1 regularization to encourage decoding generalizability and increase sensitivity for sequence detection (Liu, Dolan, et al., 2021).

To quantify the decoding accuracy, we used a ten-fold cross-validated method on the data acquired in the localizer task. The predicted accuracy was defined as the mean proportion of test trials where the classifier exhibiting the highest probability corresponded to the trial label (Fig. 3a and Extended Data Fig. 2). To test the significance of decoding accuracy and control for multiple comparisons over time, we used the nonparametric permutation test to identify the significant decoding accuracy from 0 to 750 ms post-stimulus onset.

To ensure that the decoding accuracy was equivalent across 29 states for reliable sequenceness measures, we conducted a repeated-measure analysis of variance (ANOVA) with the number of states as the independent variable and the peaked decoding accuracy (at 200 ms post-stimulus onset) as the dependent variable. Moreover, to further confirm there were no associations between the decoding accuracy and the ranks in each dimension, we used a linear mixed-effect model to fit the decoding accuracy with fixed-effect variables of ranks in attack power and defense power (Eq. 1).

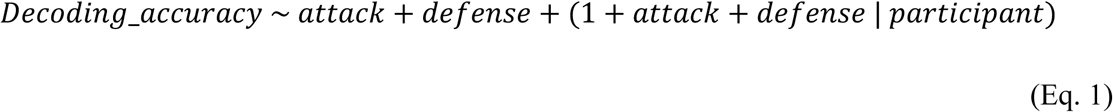

To further evaluate the temporal generalizability of our classifiers, we conducted an out-of-sample decoding analysis. Specifically, classifiers trained on all trials of stimulus-evoked neural activity from the localizer phase (i.e., the initial session) were applied to data from subsequent task phases. This training set was identical to that used for the sequenceness analysis described below. Neural responses from the learning and inference phases were segmented into epochs corresponding to stimulus presentations (e.g., “bottle” > “wallet” trials during piecemeal learning; see Extended Data Fig. 3), which served as independent test sets.

Out-of-sample decoding accuracy was calculated as the mean proportion of test trials in which the classifier correctly identified the object label. We found that classifiers generalized well to the later phases of the experiment, with peak decoding accuracies exceeding 25%, substantially above chance (see Extended Data Fig. 3 for details).

### Sequenceness measure

After determining that decoding accuracy peaked at 200 ms post-stimulus onset, we applied classifiers trained at this time point to channel-level resting or on-task MEG data, generating time series of decoded states for sequence detection (Fig. 2b). During offline rest and on-task inference, we focused on the sequential neural reactivations of task states forming 1D hierarchies for either attack or defense power, as well as 2D replay of diffusive transitions within the 2D map throughout the learning process (see Extended Data Fig. 4a). The 2D diffusive replay was further divided into central and boundary relationships, modeled as two separate regressors in the same GLM (see Extended Data Fig. 5a). Using TDLM (Liu, Dolan, et al., 2021; Liu et al., 2019; Liu, Mattar, et al., 2021; Nour et al., 2021; Wimmer et al., 2020), we quantified the strength, termed “sequencessness”, as the degree to which state representations reactivated in a hypothesized transition structure at a specific time lag (Fig. 2, c–e). We also examined how the strength of specified replay changed during the piecemeal learning and local learning phases, termed “sequenceness changed over time”.

TDLM uses a two-stage regression framework to quantify replay strengths (“sequenceness”) and its temporal change (“sequenceness changed over time”). In the first-stage model, we regress the reactivation time course of each state (*X*_*j*_, *j* ∈ [1:29]) on the time-lagged reactivation predictors of all states (*X*(Δ*t*)_*i*_, *i* ∈ [1:29]). As shown in Extended Data Fig. 7c (upper panel), we include a “reactivation” regressor, *X*(Δ*t*)_&_, capturing the main effect of replay strength at each time lag, and a “reactivation × time” regressor, *Xtime*(Δ*t*)_*i*_ (Extended Data Fig. 7c, bottom panel), to assess how replay strength changes over time. This method has been validated in previous methodological work (Liu, Dolan, et al., 2021). It will be useful to quantify learning-related change during the piecemeal learning and local learning phases. Formally,

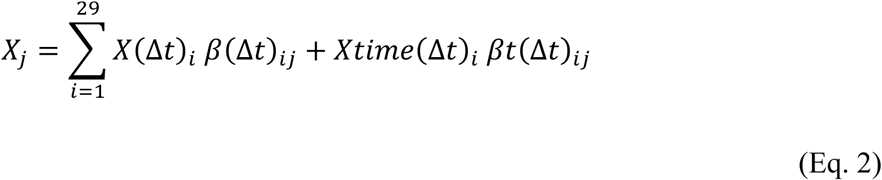

Here, *β*(Δ*t*)*_ij_*. quantifies the empirical transition from state *i* to state *j* at time lag Δ*t*, while *βt*(Δ*t*)*_ij_* indicates how that transition strength changes over learning. All these regression coefficients (main effects and time-modulated effects) form an empirical transition matrix *B* in the first-stage model.

In the second-stage model, we test for specific replay transitions of interest: 1D replay along each dimension (attack or defense) and 2D replay for an integrated map, by simultaneously fitting their hypothetical transition matrices *T*_*r*_ (including *T*_1*D*_1*st*_, *T*_1*D*_2*nd*_, *T*_2*D*_), to the lag-specific empirical transition strengths or their temporal changes (Extended Data Fig. 4a). In addition, we include an identity matrix (*T_auto_*) to model self-transitions and a constant matrix (*T_constant_*) to account for general background activity:

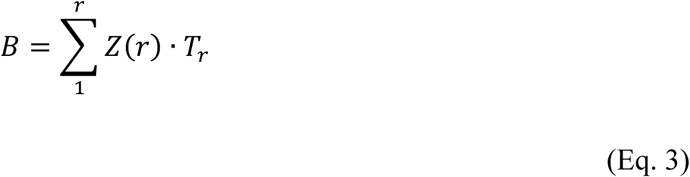

where the regression weights *Z*(*r*), quantify the replay strength or its changes over time of each replay predictor (i.e., 1D replay of the 1^st^ dimension, 1D replay of the 2^nd^ dimension, and 2D replay).

We use non-parametric permutation tests (1,000 permutations) for statistical inference. Specifically, we shuffle rows and columns of the replay transition matrices and record the peak absolute mean sequenceness (over participants and all computed time lags) in each permuted dataset. A sequenceness value in the unpermuted data is deemed significant at peak-level FWE *p <* 0.05 if it exceeds the 95^th^ percentile of these permuted peaks. This procedure controls for multiple comparisons across time lags.

Across the entire study, the replay strength was assessed with general thresholds controlling for multiple comparisons. Because the duration of reactivation time courses and the number of replay transitions varied across phases (e.g., 5-minute vs. 2-minute resting periods, boundary vs. central replays), a single universal threshold would have been too lenient for shorter segments or too stringent for longer ones. Instead, we derived an appropriate threshold for task phases with consistent duration and number of transitions, defining significance as exceeding the 95^th^ percentile of the within-permutation peak. Under these thresholds, replay strength or its temporal change was considered significant if it exceeded that 95^th^-percentile value for the respective dataset. The sequenceness values represented the degree of replay strength for learned or unexperienced relationships during offline rest and online inference tasks. Sequenceness with shorter time lags reflects greater time compression (i.e., faster replay). Positive time-modulated effects of sequenceness values indicated the replay strength with a specific time lag increased over time during rest or tasks.

To examine replay prioritization in piecemeal learning, the transition matrix of the 2D diffusive replay was modeled separately for central and boundary relationships as two regressors in a single model (Extended Data Fig. 5a). Moreover, we modeled the replay for each learning pair during resting periods within and following the piecemeal learning phase. For piecemeal learning, replay strength was also fitted with a linear mixed-effect model, including fixed-effect variables such as learning dimension, learning pair (central vs. boundary), and learning block, along with their two- and three-way interactions (Eq. 4).

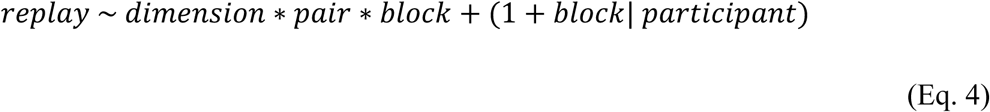

Analogous to replay detection during rest, we detected the on-task replay during inference using TDLM with some modifications. We focused on the trial-related pairs for all inference phases. The trial-related pairs during inferences (i.e., *S*1- and *S*2-related replay, Fig. 5a, d) were defined as the 2D diffusive replay related to two stimuli, including the trial-specific trajectory *S*1 – *S*2.

On-task replay detection was performed on the entire trial process rather than rest or intertrial intervals. The 2D diffusive replay were divided into trial-relevant transitions (*S*1- and *S*2-related replay, see Fig. 5a, d) and trial-irrelevant transitions (i.e., 2D diffusive replay excluding trial-related pairs). We also tested the time modulation effect of trial-related replay across all trials during inference. During rest sessions, we controlled for alpha oscillations to mitigate their potential confounding influence on sequenceness estimates (see Liu, Dolan, et al., 2021, for justification). Alpha power was not regressed out in the on-task replay analysis, as participants were actively engaged in decision-making during inference.

To assess group-level replay effects, we calculated average sequenceness for each participant across all correct-response trials. This was done separately for trial-relevant and trial-irrelevant 2D replay, in addition to 1D replay for the 1^st^ and 2^nd^ learned dimensions. Finally, to investigate the behavioral relevance of on-task replay, we used generalized linear mixed-effect logistic models to predict trial-by-trial inference accuracy based on replay strength for the corresponding inferred trajectories.

### Identification of replay onset

As significant replay was found during post-learning rest sessions (i.e., rest after piecemeal learning, before map-based inference and after local learning), we then identified the time points (i.e., replay onset) with high reactivation probability of a stimulus followed by another stimuli (e.g., surrounding states of a lagged reactivated state in 2D replay) with a specific time lag.

Given the replay onset identified by the peaked time lag across sessions, we epoched the MEG data (400 Hz) of these resting sessions around the putative replay onsets (replay probability > 95^th^ percentile). To ensure that each captured event was the onset of replay event instead of the middle of a longer sequence, we employed an additional constraint to remove the candidate replay event with a high probability of replay 200 ms before the current replay onset (replay probability < 90^th^ percentile), as in our previous studies (Liu et al., 2019; Nour et al., 2021). This applied to both fast and slow replay, regardless of whether it occurred during rest or tasks. For on-task replay, we ensured that the events for fast and slow replay did not overlap.

### Time frequency analysis

To investigate the ripple power change around the offline replay onset during rest, we first extracted the replay events with a specific time lag based on the established analysis pipeline (Liu et al., 2019; Liu, Mattar, et al., 2021; Nour et al., 2021). Specifically, replay onset was identified as the time point exhibiting high replay probability (> 95^th^ percentile of session-specific replay probability distribution in each individual). Accordingly, we also identified the on-task replay (i.e., fast and slow replay) events during the inference task. For offline replay during rest, we extracted the replay events from the three resting sessions and computed a frequency decomposition (Morlet wavelet transformation, ‘wavelet’ function in MATLAB, wave number *k*_0_ = 5) for each whole-brain channel and replay event, and finally averaged the estimates over replay events and channels. To test whether there existed a power increase around replay onset relative to pre-replay period in the ripple frequency band (120-150 Hz) as in our previous studies (Liu et al., 2019; Liu, Mattar, et al., 2021; Nour et al., 2021), we calculated the mean power increase across subjects and used a nonparametric test for multiple comparisons at the cluster level. The same procedure was applied to on-task replay during inference.

### Source localization

Having confirmed that ripple power increased at replay onset, we then explored the neural source associated with this power increase in the brain. First, the individual structural MRI image was co-registered with the MEG data through the alignment of the set of fiducial markers with the preauricular points and the nasion. The co-registered subject-specific MRI was then segmented. Forward models were generated based on a single shell using superposition of basis functions that mapped the plane tangential to the MEG sensor array. Linearly constrained minimum variance beamforming (Van Veen et al., 1997) was employed to reconstruct the epoched MEG data of replay events to a grid in the MNI space with a grid step size of 5 mm. No temporal smoothing or average was implemented.

For offline resting periods, as in our previous studies (Liu et al., 2019; Nour et al., 2021), the MEG data restricted to 120-150 Hz were used to estimate the sensor covariance matrix for beamforming. After rejecting artifactual epochs, all replay events were baseline-corrected at the source level. The whole-brain voxels were estimated by subject-specific ripple power in the frequency band of 120-150 Hz. During inference, for unbiased comparison, we targeted on a broader ripple band, also termed “high gamma” (80-180 Hz) on the epoched MEG data of fast and slow on-task replay events. For both events, the subject-level gamma band power in the whole-brain voxels was estimated. The fast- and slow-replay-specific neural activity was calculated by contrasting the whole-brain estimates of fast vs. slow on-task replay. At the group level, nonparametric permutation tests were employed to identify significant clusters across the whole brain (*p <* 0.05, whole-brain FWE corrected, cluster-defining threshold: *t* > 3.1, 5000 permutations).

To target the mental navigation on a 2D map, similar pipeline was applied to identify the neural sources induced by map-based inference. Different from the procedure for replay events during rest, we epoched the navigation-related events during 2D inference (i.e., from 500 ms before *S*2 onset to the end of decision phase). The MEG data were restricted to 4-8 Hz for detections of grid-like representations during mental navigation (D. Chen et al., 2018, 2021; Maidenbaum et al., 2018). To test the specificity of hexagonal grid-like code in the entorhinal cortex and the medial prefrontal cortex, we used binary masks to extract the trial-by-trial source-level neural activities from regions of interest (ROIs) listed above and some regions of no interest as control regions such as the primary visual cortex (V1) and the primary motor cortex (Glasser et al., 2016), and tested the neural activities in other frequency band. The output data from these regions were then used as input in grid-like coding analysis (see Grid-like coding below).

### Grid-like coding

During 2D inference, transitioning from stimulus 1 (*S*1) to stimulus 2 (*S*2) can be viewed as “traversing” a trajectory on the mental 2D map. We tested whether these navigating trajectories exhibit six-fold, hexagonal grid-like coding (Bellmund et al., 2016), focusing on theta-band (4–8 Hz) source-level activity in the EC and mPFC. To rule out spurious findings, we repeated the procedure in control regions (e.g., primary visual and motor cortices) and in other frequency bands (delta: 1–4 Hz, alpha: 8–12 Hz, beta: 12–30 Hz, low gamma: 30–80 Hz, high gamma: 80–180 Hz), none of which showed a significant six-fold pattern (Extended Data Fig. 9).

We applied an RSA-based approach (Diedrichsen & Kriegeskorte, 2017; Walther et al., 2016) from *S*2 onset until the end of decision (Fig. 6a). First, trial-by-trial source-level MEG data from the EC and mPFC (Glasser et al., 2016) were baseline-corrected and averaged across voxels within the theta band. We then computed correlation distances (1 – *r*) between each pair of trajectory angles to form an empirical representational dissimilarity matrix (RDM) for each time point. Next, we built a theoretical RDM reflecting a six-fold hexagonal structure: given the six-fold periodicity (spanning the entire circle with recurring periods of 60°), pairwise trajectory angles ∼30° apart (15°–45°) were set to 1 (high dissimilarity), while others (< 15° or > 45°) were set to 0. As controls, we also tested four-, five-, seven-, and eight-fold symmetries, none of which yielded significant effects. Finally, we regressed the empirical RDM against the theoretical RDM in a second-stage GLM to estimate grid-like strength from *S*2 onset. Nonparametric cluster-level FWE corrections (Eldar et al., 2020) identified significant time windows of grid-like coding in theta-band activity (Extended Data Fig. 9). The six-fold effect emerged solely in the EC and mPFC, aligning with a recent human iEEG study that recorded directly from these regions and confirmed a grid-like code, also in theta band, for representing a 2D conceptual space (Xiao et al., 2025).

### Control analyses for head position and orientation

Given the length of the scanning session, a potential concern was that participant fatigue or head movement might compromise data quality. To mitigate this, we incorporated brief breaks after each run and longer breaks between task sessions. In addition, we performed post hoc control analyses to assess whether head motion may have influenced the neural results.

Although our MEG acquisition system (version 5.0.2) did not support continuous tracking of head position or orientation throughout the session, we recorded the initial head localization data at the start of each scanning run. We correlated these measures – i.e., the relative head position and orientation within each task session – with key neural variables, including offline and online replay strength as well as grid-like map representations. No significant associations were found between head position/orientation and any of these neural measures (see Extended Data Fig. 11). These findings suggest that head motion did not systematically bias the results and is unlikely to explain the observed neural effects.

### Statistical analysis and software

MEG preprocessing, time-frequency analysis, source localization were all performed using MATLAB R2019b (https://www.mathworks.com/) with functions from the OHBA Software Library (OSL, https://github.com/OHBA-analysis/osl-core/), Statistical Parametric Mapping (https://www.fil.ion.ucl.ac.uk/spm/software/spm12/), FMRIB Software Library (FSL, https://fsl.fmrib.ox.ac.uk/fsl/fslwiki/) and FieldTrip (https://www.fieldtriptoolbox.org). The replay analysis, RSA and grid-like coding analyses were conducted with custom MATLAB code.

The descriptive statistics were reported as mean ± 1 standard error of the mean (Mean ± *SE*) across the entire study. Statistical analyses on mean difference significance tests (i.e., *t*-test and ANOVA) were performed with a significance threshold of two-tailed *p <* 0.05. We also controlled for multiple comparisons using either false discovery rate (FDR) correction or family-wise error (FWE) correction. To compare the behavioral performance and replay for boundary and central relationships during piecemeal learning, we used the bootstrapped sampling methods to estimate the subject-level mean for each condition and the mean difference between boundary and central pairs, indicated by the 95% confidence intervals. Using bootstrapped sampling methods ensured unbiased estimation across conditions with different number of trials. All individual-level correlational analyses (e.g., neural replay and behavioral performance) employed Spearman correlation analysis across the study. To further increase the robustness of these findings, significance tests were repeated after excluding outliers (data points beyond 3 standard deviations from the mean), if any were present. Besides, linear mixed-effect models for decoding accuracy across states were applied on the state-specific data in each participant and were estimated using restricted maximum likelihood using lme4 and lmerTest in R (Bates et al., 2015; Kuznetsova et al., 2017). We used the Type II Wald chi-square tests to identify the significance of main effects and interaction effects. Generalized linear mixed-effect models for learning performance and for nonlocal inference accuracy were performed on trial-wise data and fitted using maximum likelihood (Laplace approximation). For nonparametric permutation test in sequenceness measure, the time lags corresponding to the sequenceness or time-modulated sequenceness changes in the unpermuted data surviving the 95^th^ percentile of the null distribution obtained from 1000 permutations was deem significant (*p <* 0.05, peak-level FWE corrected). For the RSA and grid-like coding analysis, as well as the associations between offline/on-task replays and the grid-like code across time, the time window in the unpermuted data with its sum-of-*t* value surviving the 95^th^ percentile of the null distribution was determined significant (*p <* 0.05, cluster-level FWE corrected) (Eldar et al., 2018).

### Visualization

For replay analyses, we used the decoded time series of state reactivations during both offline rest and on-task inference. To illustrate these replay events, we plotted the empirical time courses of various task states around the replay onset (see Fig. 2b, Fig. 5c, and Fig. 7). Vertical bars in these plots indicate any state reactivation probability exceeding the 95% threshold of overall probabilities across all states. The shaded interval from replay onset to a specified time lag highlights the replay event.

To generate the behavioral and neural replay heatmaps for each state in piecemeal learning (Fig. 3b, d), we first computed the mean test accuracy and replay strength per state for each participant, based on the learning pairs or 2D diffusive transitions that included that state. We then averaged these values across participants and normalized them to a 0–1 range for visualization. This normalization does not affect the statistical conclusions. In fact, single GLMs that model boundary and central replays simultaneously, along with bootstrap analyses, confirm that the observed differences remain robust, regardless of the number of transitions or presentations for each state.

## Data and code availability

MEG and behavioral data that supporting the conclusions in this study will be available on https://osf.io/xvpuz/ upon publication. The analysis code will be publicly available on https://gitlab.com/liu_lab/form_2dmap.git upon publication.

## Acknowledgments

We thank Kristopher Jensen, Mathias Sablé-Meyer, Sebastijan Veselic, Alon Baram, Eleanor Spens, and Avital Hahamy for their critical reading and comments on the initial manuscript. We are deeply grateful to all participants for their generous time and dedication to this extended study.

## Author contributions

Conceptualization: YL, JO, YQ, TB

Methodology: JO, YL, TB

Investigation: JO, YX, ZX, YL

Visualization: JO, YL

Funding acquisition: YL

Project administration: YL

Supervision: YL

Writing – original draft: JO, YL

Writing – review & editing: JO, YL, TB

## Extended Data Figures

**Extended Data Fig. 1.**
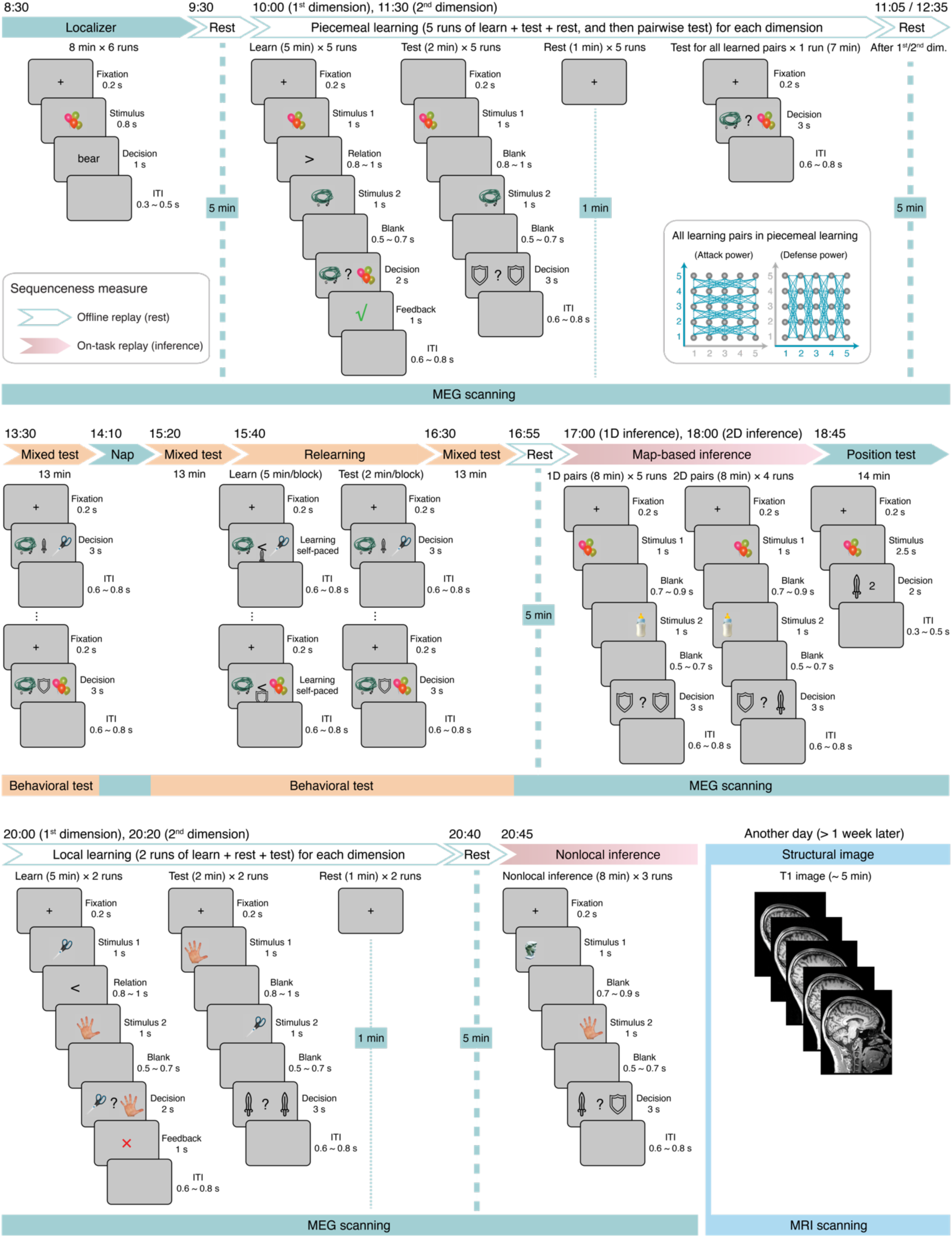
Experiment schedule and core trial structure. (**Overall design**) All participants followed a similar experimental schedule and trial structure, with only minor variations in timing. Each full-day MEG session lasted approximately 13 hours, including roughly 8 hours of scanning. To mitigate fatigue, each MEG run was limited to ∼8 minutes, after which participants took brief breaks. Every hour, they had a 15–20-minute extended break, during which they could exit the scanner. At the start of the experiment, each subject’s head position was marked (in red pen) and monitored throughout; participants were asked to return to this initial position after each break (see Methods for details). Post-experiment analyses confirmed stable head positions with no correlation between head motion and key task measures like replay and grid-like representation (Extended Data Fig. 11). The figure below illustrates a typical experimental schedule and trial structure. (**Upper panel: morning session, ∼8:30 a.m.**) Participants began with the localizer phase, where they viewed an object picture and decided whether the accompanying text matched it (Fig. 1b). Immediately following, they underwent a 5-minute rest scan (eyes open, no task). Around 10:00 a.m., they commenced the piecemeal learning phase, learning object pairs that differed by one rank in the target dimension (attack power or defense power). Each feature dimension was learned over five runs, each containing 20 unique pairs (presented twice, e.g., “balloon” > “hose” and “hose” < “balloon”), giving 40 learning trials plus 20 testing trials (no feedback), followed by a 1-minute rest. After each dimension, participants were quizzed on all 100 pairs, and then took a 5-minute rest. All one-rank-difference pairs within the dimension are depicted using green, bidirectional arrows (bottom right). (**Middle panel: afternoon session, ∼1:30 p.m.**) After completing the learning phases, participants took an intermixed test on the two dimensions outside the scanner, then had a nap in the scanner. Around 3:20 p.m., they performed an intermixed test. Next, they relearned any incorrectly answered pairs through two rounds of learning + testing outside the scanner, followed by another intermixed test. Once these relearning phases were done, participants returned for MEG scanning and had a 5-minute rest before starting the map-based inference phase (∼5:00 p.m.). It contained two tasks: 1D inference (two objects presented, participants identified the higher rank within one dimension) and 2D inference (participants compare whether one object’s attack power exceeded another’s defense power). Participants then completed the position test to show their understanding of the 2D map space. (**Bottom panel: evening session, ∼8:00 p.m.**) Four new objects were introduced to the central area of the map. Participants engaged in local learning to establish rank relationships between each new object and its neighbors in each dimension. Following a 5-minute rest, they performed nonlocal inference, inferring relationships between the new objects and non-adjacent objects never directly learned — thus “nonlocal.” One week later, participants returned for an MRI session to acquire T1-weighted structural images for source-localization analysis of the MEG data. In each panel, task names shaded in green indicate phases with MEG scanning; names shaded in orange represent phases with only behavioral recordings. Phases framed in blue-green were used for offline replay analyses (rest periods), while those shaded in sangria-pink denote on-task replay analyses.

**Extended Data Fig. 2.**
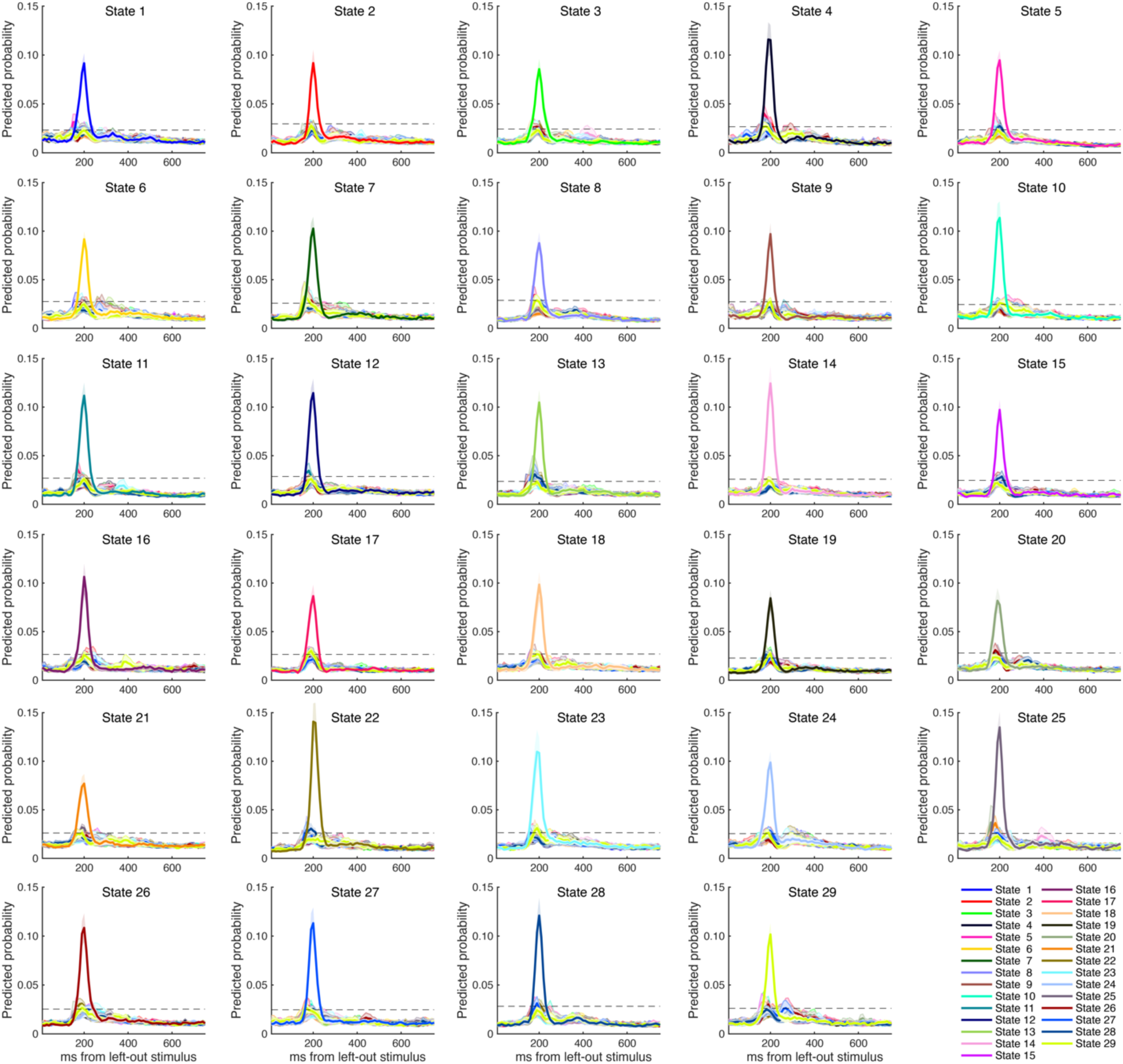
Classifier performance for each of 29 states. Using lasso regression models in the localizer task, we obtained the ten-fold cross-validated predicted probability for each classifier. These classifiers were trained at 200 ms post-stimulus onset and subsequently tested on all states at different time points (represented on the *x*-axis). Line colors indicate different states. The predicted accuracy notably increases when the target state aligns with the classifier state, reaching its peak at 200 ms after state onset. The dashed line represents the permutation threshold, and the shaded area reflects the *S.E*.

**Extended Data Fig. 3.**
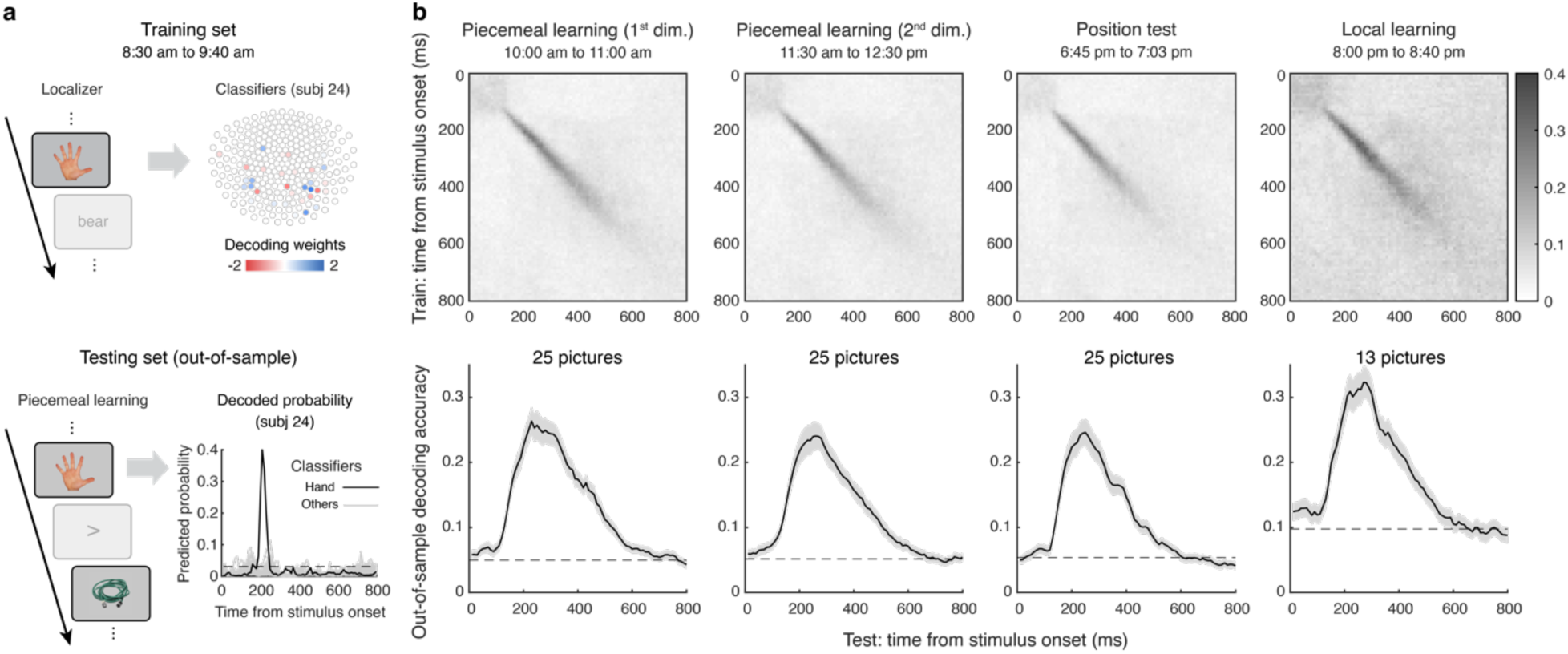
Out-of-sample generalization of classifiers across task phases. **a,** Illustration of the out-of-sample generalization procedure. Classifiers were trained using stimulus-evoked neural data from the localizer phase (training set; see top panel for empirical beta weights of the “hand” classifier in subject 24). To assess generalization, these classifiers were applied to neural data from object presentation periods in later task phases (testing set). Predicted probabilities were significantly higher when the classifier matched the presented object, as shown in an example from subject 24 (bottom panel). **b,** Mean out-of-sample decoding probabilities across subsequent task phases. Classifier performance was highest when training and testing occurred at the same time point after stimulus onset (top panel). Peak decoding accuracy occurred around 200 ms post-stimulus onset (bottom panel) and was consistent across task phases, indicating robust generalizability of the classifiers over time without significant time-dependent degradation.

**Extended Data Fig. 4.**
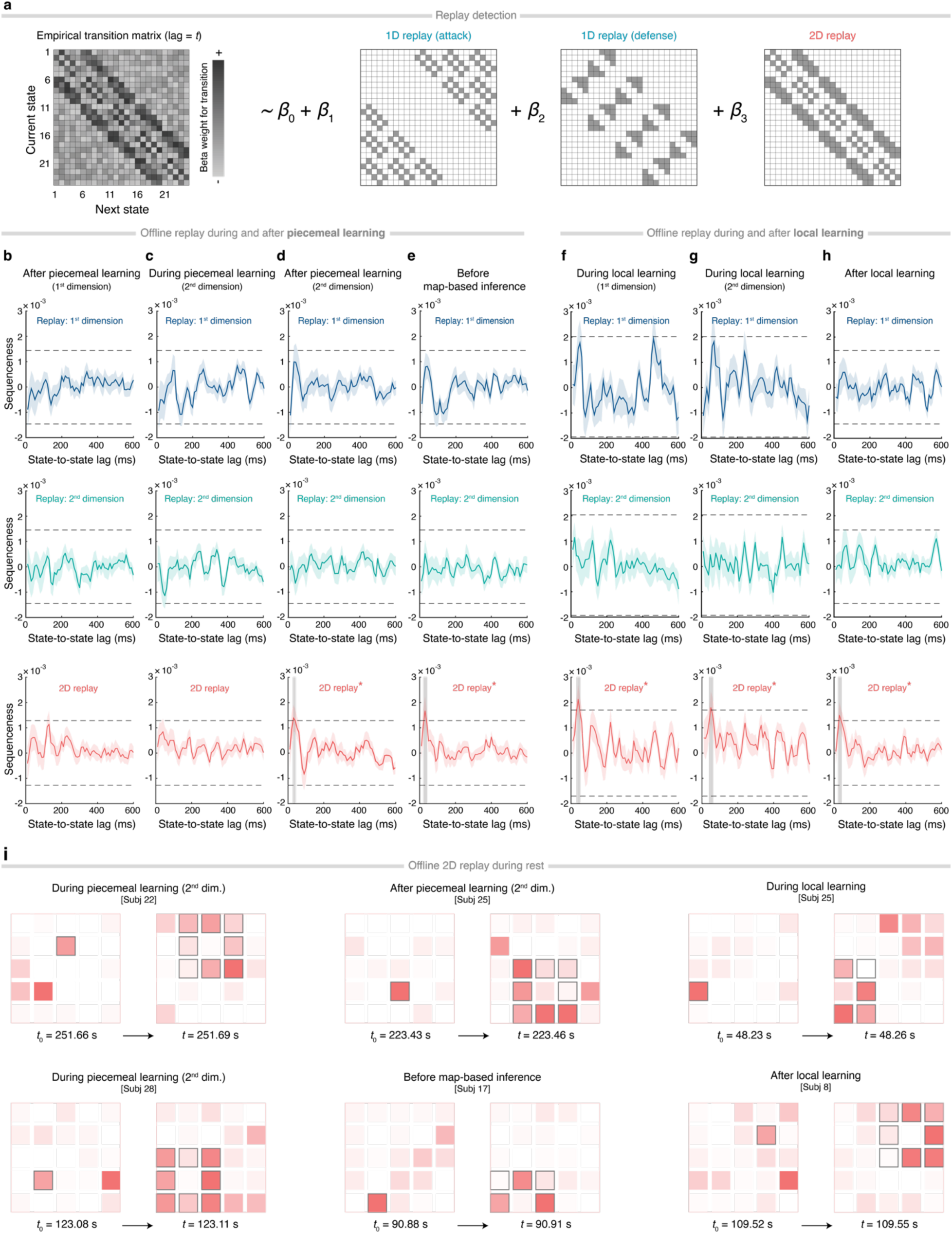
Offline replay during and after learning. **a**, Replay detection using Temporally Delayed Linear Modeling (TDLM). State-by-state matrices for 1D and 2D replay were modeled simultaneously as predictors to capture the sequential reactivation strength. **b–f**, Offline replay strength separately for 1D hierarchy, measured during (**c**) and after piecemeal learning (**b**, **d**, **e**) as well as during (**f**, **g**) and after local learning (**h**). Across sessions, neither of the 1D replays survived the permutation threshold (dashed line). No significant evidence was observed for 1D replay of either 1^st^ or 2^nd^ dimension across the whole experiment. However, 2D replay was found immediately after piecemeal learning and persisted thereafter. **i**, These additional examples of 2D replay, drawn from different participants, are less pronounced than the illustrative case in Fig. 3d but still exhibit identifiable 2D replay features. They highlight the variability in individual data while confirming that, at the group level, 2D replay remains robust and statistically reliable. Shaded regions around the mean line indicate the *S.E*. The dashed line represents the general permutation threshold for each specified replay across the experiment. The gray shaded time window in the replay graphs highlights the time lags surviving permutation. \**p <* 0.05.

**Extended Data Fig. 5.**
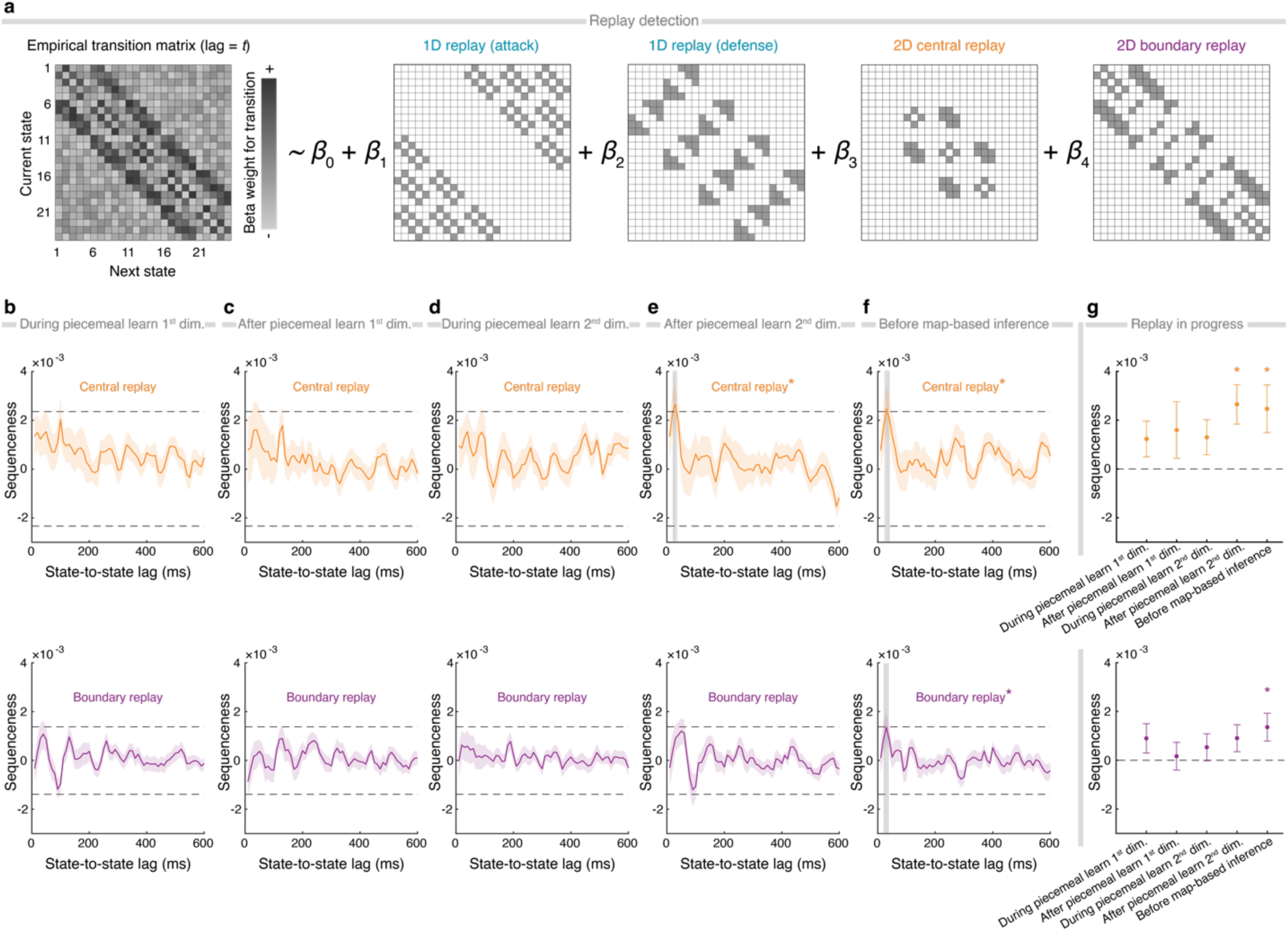
Offline 2D replay over the central and boundary relationships during and after piecemeal learning. **a**, Replay detection using Temporally Delayed Linear Modeling (TDLM). State-by-state matrices for 2D replay were modeled separately for central and boundary pairs as two predictors simultaneously to capture the sequential reactivation strength. This effect was estimated independently for each time lag (*t*) and the FWE-corrected significance threshold was defined by a nonparametric permutation test (*N* = 1000). **b–f**, Offline replay strength of 2D diffusive transitions of central and boundary pairs measured during piecemeal learning (**b**, **f**) and after piecemeal learning (**c**, **e**, **f**). Across sessions, central replay did not survive the permutation threshold (dashed line) until the end of piecemeal learning the 2^nd^ dimension (**g**, see also Fig. 4c, left) and remained stable thereafter (**e** and **f**, gray shaded time window, peaking at a 30-ms time lag). However, evidence of boundary replay was shown on the bottom panels, it was only present before map-based inference, peaked at the same time lag as replay of central pairs. **g**, Summary of the 30-ms lag offline 2D replay across the task stages. Error bars and shaded areas indicate the *S.E*. The dashed line represents the general permutation threshold for each replay. The gray shaded time window in the replay graphs highlights the time lags surviving permutation. \**p <* 0.05.

**Extended Data Fig. 6.**
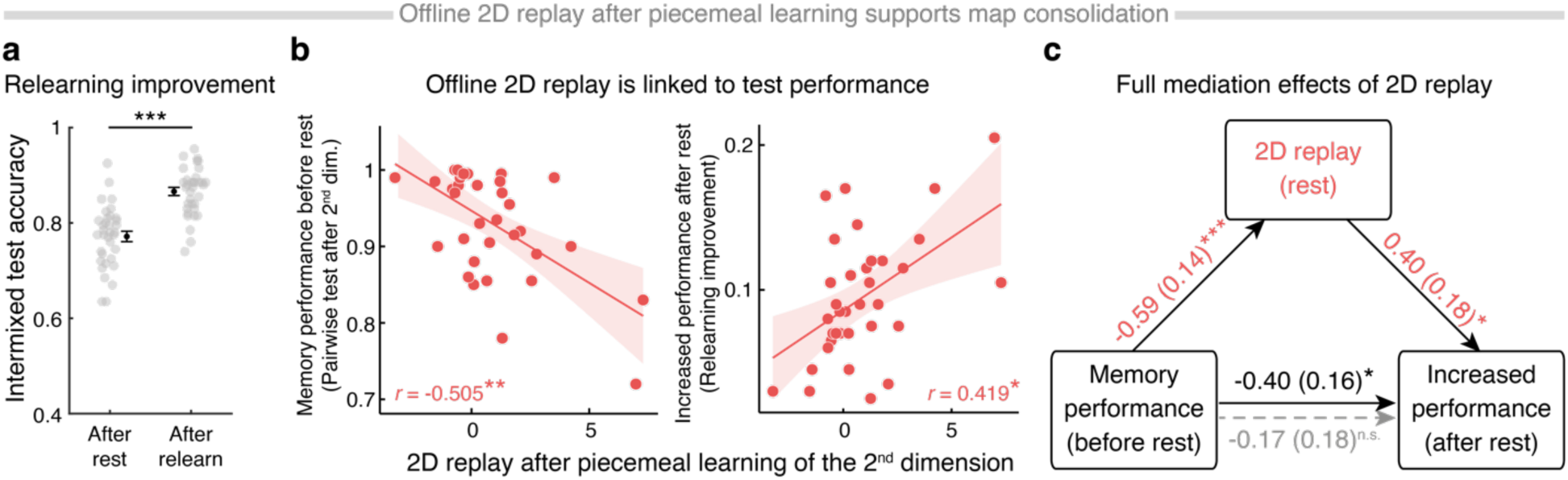
Offline 2D replay after piecemeal learning predicts map consolidation. **a**, Immediately following the post-piecemeal-learning rest, participants underwent an intermixed test where all learned pairs from both dimensions were presented in random order, assessing their knowledge of all pairs without feedback. This is termed “after-rest” test. Then, after relearning any previously incorrectly-responded pairs, participants took an intermixed test again, this is termed “after-relearning” test. A significant rise in test accuracy between these two tests reflects “relearning improvement”. **b**, The 2D replay measured during rest after piecemeal learning was negatively correlated with the memory performance before the rest period (left panel), yet positively correlated with the subsequent performance improvement after relearning (right panel). **c**, Mediation analysis indicates that 2D replay fully mediates the relationship between pre-rest memory performance and post-relearning improvement. Each point represents an individual participant. Error bars indicate the *S.E*. The shaded area of the fitted lines in the correlation plots depicts the 95% confidence interval (CI). \**p <* 0.05, \*\**p <* 0.01, \*\*\**p <* 0.001, *n.s*.: non-significant.

**Extended Data Fig. 7.**
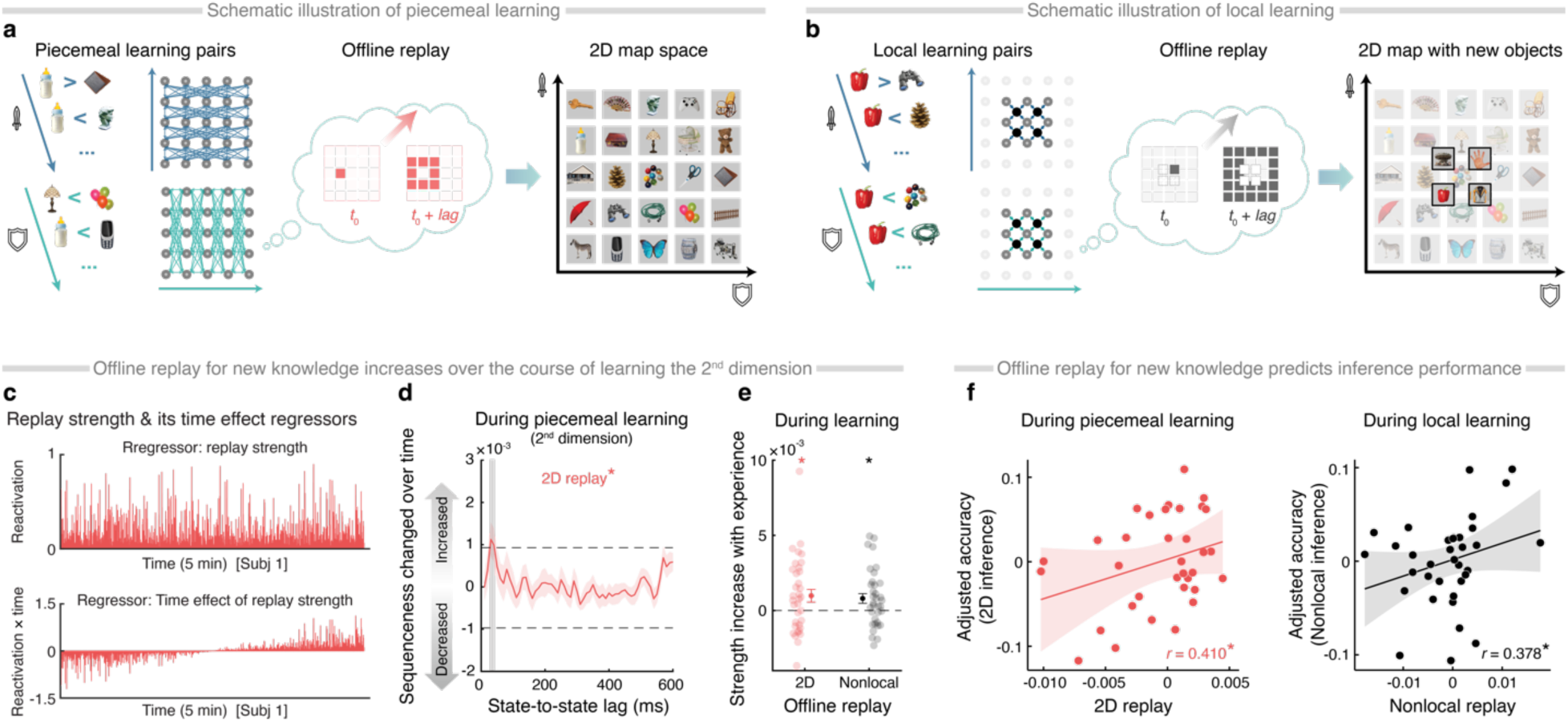
Offline replay during learning predicts novel inference performance. **a**, A schematic illustration of piecemeal learning, in which participants learned pairwise rank relationships (one-rank differences) among 25 objects in each dimension. Offline 2D replay increased over the course of piecemeal learning of the 2^nd^ dimension, facilitating the development of an integrated 2D map representation. **b**, A schematic illustration of local learning, during which four new objects were introduced into the central area of the existing 5×5 map. Similar to piecemeal learning, participants learned rank relationships in each dimension, but only for the new objects and their adjacent neighbors. During local learning of the 2^nd^ dimension, “nonlocal replay” of newly acquired knowledge (i.e., relationships between new objects and non-neighboring objects) increased, reflecting the construction of a “new” map. **c**, To capture both the main effect of specific replay strength (upper panel) and its time modulation (lower panel), two different regressors are used in the first-stage GLM. Specifically, the “reactivation” regressor (upper) is a lagged copy of each task state’s reactivation strength, while the “reactivation × time” regressor (lower) is the element-wise product of state reactivation and z-scored time. This method has been validated in Liu et al. (2021). **d**, The “sequenceness changed over time” reveals that 2D replay strength at the 30-ms lag (peak) increases significantly during piecemeal learning of the 2^nd^ dimension, after controlling for multiple comparisons. **e**, Replay strength increased significantly over the course of the 2^nd^ dimension for both 2D replays in piecemeal learning and nonlocal replays in local learning. These forms of offline replay appear to complement learning of the task representation. **f**, Offline replays from the 2^nd^ dimension in both learning phases positively predict the subsequent inference performance, even when controlling for learning performance. The left-panel result remained significant after excluding one outlier (*r* = 0.372, *p =* 0.033). Each dot represents an individual data point. The shaded area around the mean line and error bars indicate the *S.E*. The shaded area of the fitted lines in the correlation plots depicts the 95% confidence interval (CI). \**p <* 0.05.

**Extended Data Fig. 8.**
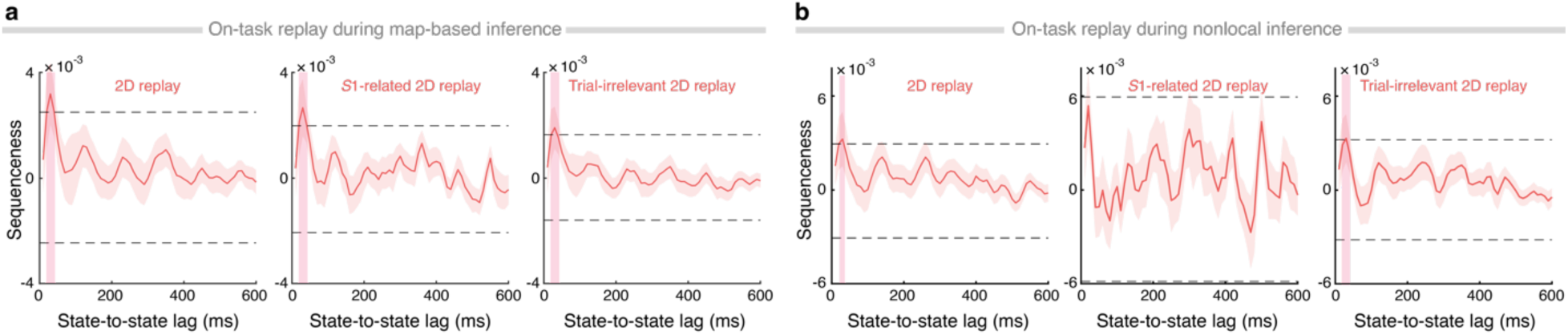
General on-task fast replays during inference tasks. For both (**a**) map-based inference and (**b**) nonlocal inference, 2D diffusive replays (for both trial-relevant and trial-irrelevant pairs) were also detected on-task, all at a fast replay speed (significant replay speed range highlighted in pink, peaking at a 30-ms time lag). For trial-relevant pairs, the *S*1-related diffusive replay was evident in the map-based inference task. A similar trend was observed in the nonlocal inference phase. Additionally, the 2D diffusive replay of trial-irrelevant pairs (excluding *S*1- and *S*2-related) was significant for all inference sessions. Shaded areas of the mean line represent *S.E*. The dashed line represents the permutation threshold. The color-shaded time window in the replay graphs highlights the time lags surviving permutation.

**Extended Data Fig. 9.**
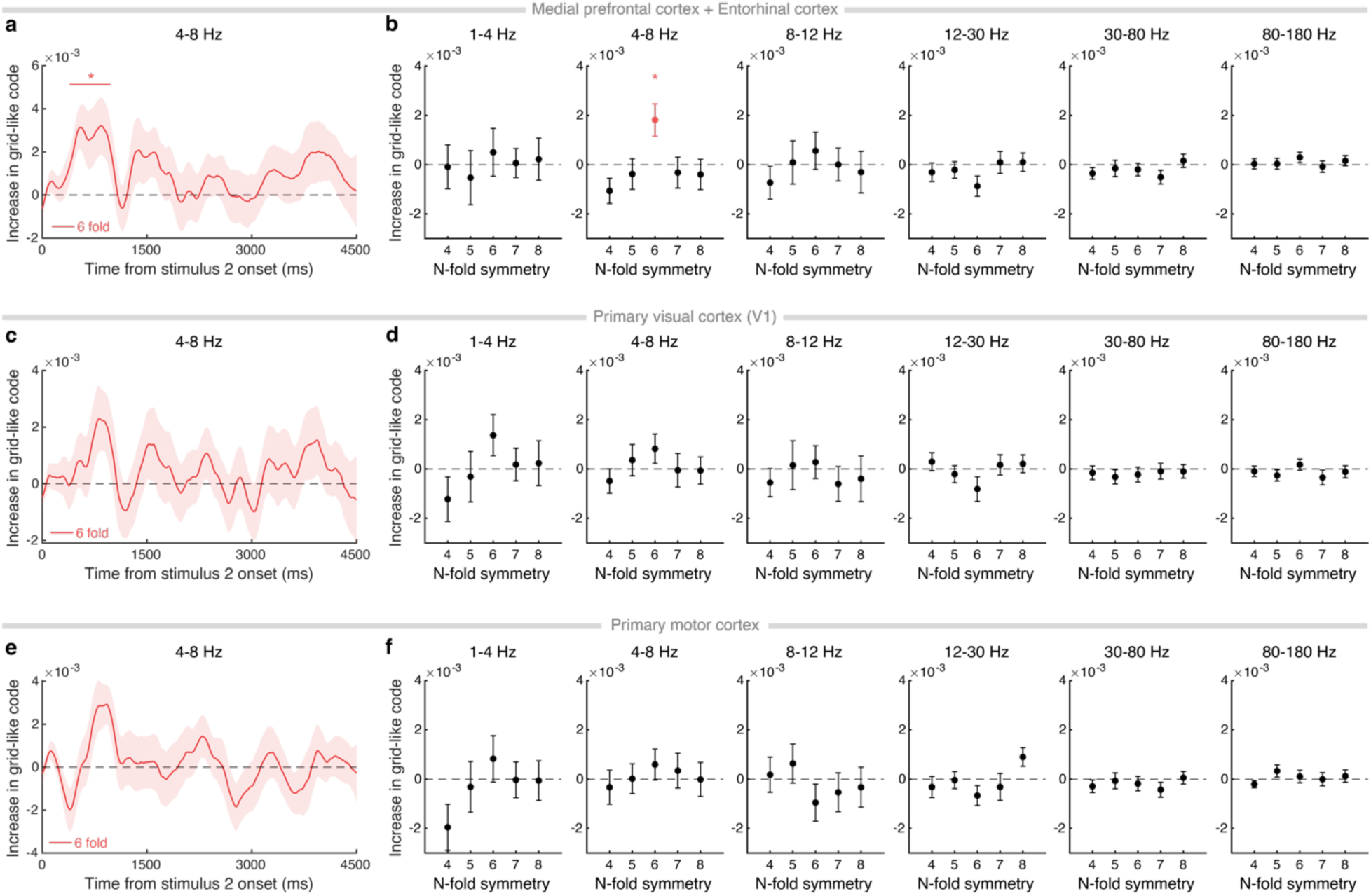
Grid-like code in all ROIs during map-based inference. **a**, **c**, **e**, We tested the specificity of the six-fold grid-like code in theta-band (4–8 Hz) activity from (**a**) mPFC and EC, (**c**) primary visual cortex, and (**e**) primary motor cortex starting from *S*2 onset. These ROIs were defined using the Human Connectome Project multimodal parcellation atlas (Glasser et al., 2016). **a**, Only the mPFC and EC showed a six-fold grid-like code, significant from 390 ms to 980 ms after *S*2 onset, cluster-level FWE-corrected. **b**, This effect was specific to the six-fold symmetry in the theta band, FWE corrected. **d**, **f**, No significant grid-like pattern emerged in the control ROIs or in other frequency bands (delta: 1–4 Hz, alpha: 8–12 Hz, beta: 12–30 Hz, low gamma: 30–80 Hz, high gamma: 80–180 Hz). Shaded regions and error bars denote the *S.E*. \**p <* 0.05.

**Extended Data Fig. 10.**
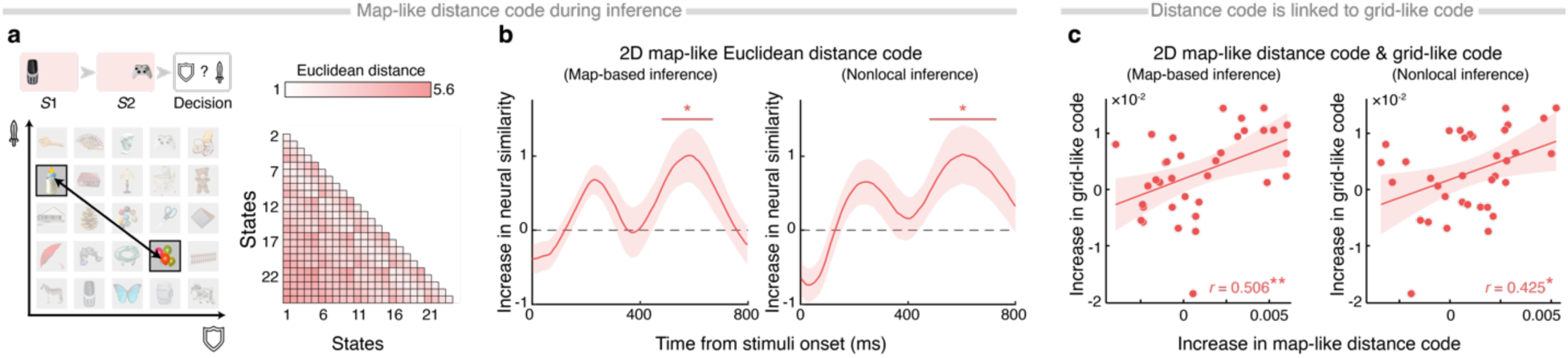
2D map-like representations measured by “distance code” during inference phases. **a**, Illustration of the distance code and the empirical RDM. The distance code reflects how neural similarity aligns with the Euclidean distance of each object in a 2D map. Such codes have been observed in the hippocampus and mPFC (Boorman et al., 2021; Park et al., 2021a). **b**, Here, the distance code was significant during both map-based inference (480–670 ms, peaking at 590 ms post-stimulus) and nonlocal inference (480–730 ms, peaking at 600 ms post-stimulus), surviving cluster-level FWE correction (*p <* 0.05). **c**, There were significant positive correlations between the Euclidean-distance codes and the EC and mPFC grid-like code (Fig. 6b, peaking at 850 ms after *S*2 onset) across both inference tasks (map-based inference: *r* = 0.506, *p =* 0.004; nonlocal inference: *r* = 0.425, *p =* 0.023; FWE corrected). Shaded regions around the mean line denote the *S.E*. The shaded area of the fitted lines in the correlation plots depicts the 95% confidence interval (CI). \**p <* 0.05, \*\**p <* 0.01.

**Extended Data Fig. 11.**
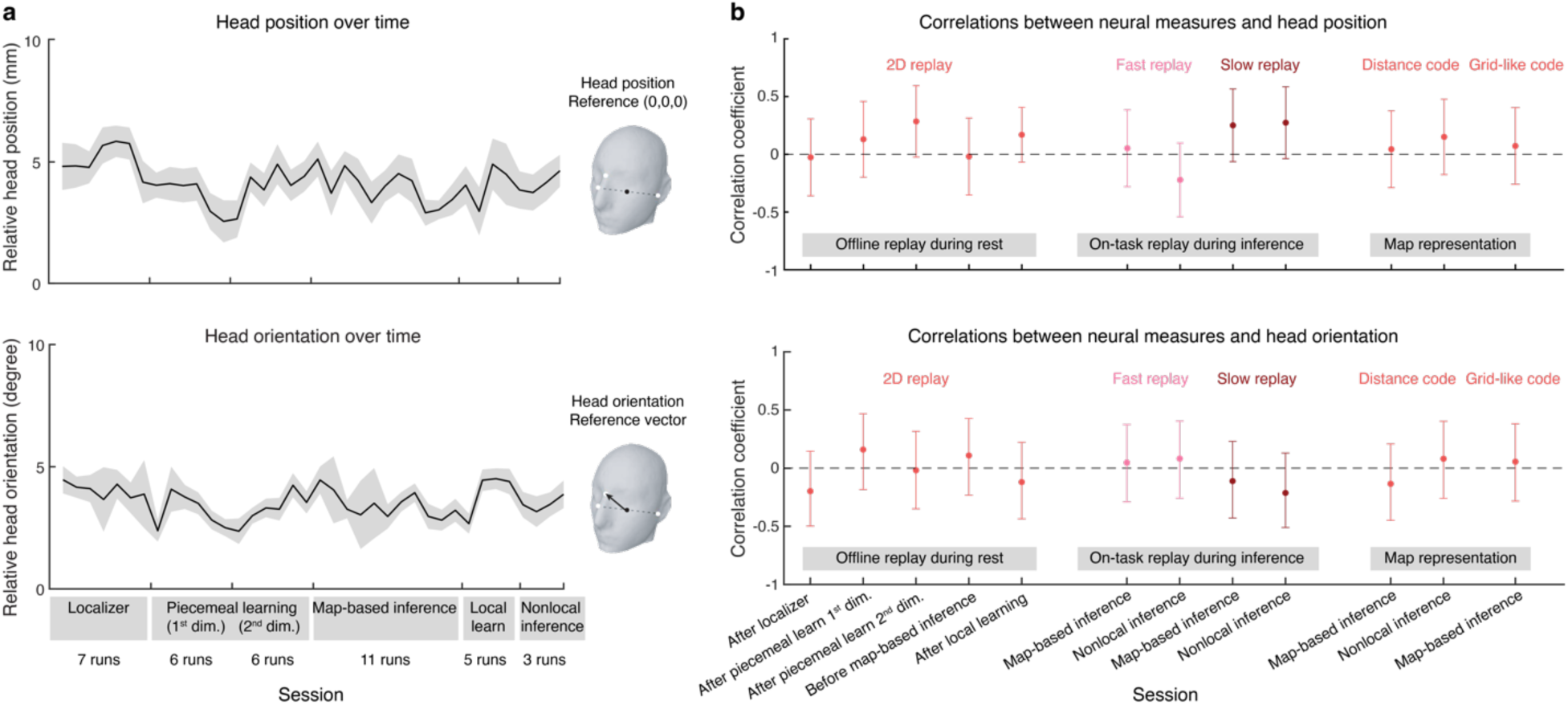
No association between head positions/orientations and key task-related measures (including neural replay and cognitive map representations). **a,** Relative head positions across task sessions. Neither head position (top panel) nor head orientation (bottom panel) showed systematic changes over time relative to each participant’s reference point (black dot) or orientation vector (black arrow), indicating stable positioning throughout the scanning session. **b,** Head position (top panel) and orientation (bottom panel) were not significantly correlated with any key neural measures, including offline replay during rest, on-task replay during inference, or grid-like cognitive map representations. Shaded areas indicate the standard error of the median. Error bars denote the 95% confidence intervals of correlation coefficients; all *p*-values > 0.1. The color scheme matches that used in the main figures.

## Supplementary Information

### Offline 2D replay prioritized weakly-learned information for map consolidation

During the 2^nd^ dimension in the piecemeal learning phase, participants performed better on boundary relationships than on central ones. Yet, an opposite pattern emerged in the offline replay: we observed stronger replay for central transitions rather than boundary transitions. One plausible explanation is that offline replay targets weakly learned information to supplement online learning (Schapiro et al., 2018). Supporting this, a significant negative correlation emerged between overall learning performance and the strength of 2D replay during the subsequent rest period [Spearman correlation: *r* = -0.505, *p =* 0.002, Extended Data Fig. 6b, left]. In other words, participants with lower initial performance exhibited stronger offline 2D replay during subsequent rest, indicating that replay generally prioritizes less well-learned relationships in the map.

Following this post-piecemeal-learning rest period, participants took an intermixed test covering all pairs in both dimensions, then relearned any previously incorrectly-answered pairs (see Methods and Extended Data Fig. 1). As expected, performance significantly improved after relearning [immediately after rest: 0.772 ± 0.011; after relearning: 0.866 ± 0.008; *t*(34) = 12.837, *p <* 0.001; Extended Data Fig. 6a]. Notably, the strength of 2D replay during rest also positively correlated with subsequent relearning improvement (*r* = 0.419, *p =* 0.012; Extended Data Fig. 6b, right), suggesting that replay does more than merely tracking previously poorly-learned pairs; it actively facilitates their consolidation and performance gains.

To test for mediation effects of offline replay, we treated pre-rest memory performance as the independent variable and relearning improvement as the dependent variable, with offline 2D replay serving as the mediator. Using the “lavaan” package in R (Rosseel, 2012), we found that the 2D replay across the entire map during post-piecemeal-learning rest fully mediated the negative relationship between pre-rest performance and post-rest relearning improvement. Specifically, participants with lower pre-rest memory showed stronger 2D replay during rest, which in turn predicted larger gains after relearning [before mediation: *c* = -0.403 ± 0.159, *p =* 0.016; after mediation: *c’* = -0.169 ± 0.179, *p =* 0.345, see Extended Data Fig. 6c]. Together, these findings indicate that offline replay during rest prioritizes weakly learned information, thereby facilitating consolidation of the 2D map.

### On-task 2D map-like representation during inference phases

During the inference phase, we observed a grid-like code in the EC and mPFC. In addition, we identified a “distance code”, capturing how neural activity patterns reflect the pairwise Euclidean distances among objects in a 2D conceptual space (Theves et al., 2019). In other words, objects that are closer in 2D Euclidean space tend to have more similar neural activation patterns (Morgan et al., 2011; Nielson et al., 2015; Park et al., 2021a; Theves et al., 2019; Viganò et al., 2021).

To detect this distance code using MEG, we employed RSA-based approach (Diedrichsen & Kriegeskorte, 2017; Walther et al., 2016), comparing neural patterns during the inference phases to those measured in a pre-learning localizer phase. First, a GLM was used to estimate state-specific neural activations (Extended Data Fig. 10a, left). We then constructed an empirical representational dissimilarity matrix (RDM) based on Pearson correlation distances (1 – *r*) between states (Deuker et al., 2016), quantifying how these representations changed from the localizer to the inference phases. A Gaussian kernel was applied to smooth the representation changes over time. Following earlier work (Park et al., 2021a), we used the Euclidean distance between each pair of states as our theoretical RDM (Extended Data Fig. 10a, right), then regressed it against the empirical RDM to derive the strength of the “distance code”.

Finally, group-level nonparametric tests identified significant time windows while correcting for multiple comparisons (Eldar et al., 2018). A robust distance code emerged during both map-based inference (480–670 ms, peaking at 590 ms, cluster-level FWE-corrected, *p =* 0.031) and nonlocal inference (480–730 ms, peaking at 600 ms, cluster-level FWE-corrected, *p =* 0.017) after stimulus onset (Extended Data Fig. 10b). The temporal evolution of this distance code reflects ongoing mental representations of the conceptual relationships, and its peak marks the moment when neural activity aligns most closely with the Euclidean structure of the 2D map.

Moreover, we hypothesized that the grid-like code, which acts as a generalizable schema, might correlate with this distance code in both the existing map (map-based inference) and the new map (nonlocal inference). Indeed, the peak strength of the distance code in both tasks was significantly correlated with the grid-like code in the EC and mPFC [map-based inference: *r* = 0.506, *p =* 0.004, FWE corrected; nonlocal inference: *r* = 0.425, *p =* 0.023, FWE corrected; Extended Data Fig. 10c]. These results suggest that grid-like coding serves as a schema representation, supporting both existing and newly learned 2D maps during inference.

